# Distinct signaling mechanisms and proteome phenotypes are elicited by compartment-specific genetic defects of copper homeostasis

**DOI:** 10.64898/2026.01.13.699066

**Authors:** Alicia R. Lane, Nadia Gonzalez, Avanti Gokhale, Stephanie A. Zlatic, Brooke M. Allen, Noah E. Scher, Anne M. Roberts, Duc M. Duong, Blaine R. Roberts, Alysia D. Vrailas-Mortimer, Erica Werner, Victor Faundez

**Affiliations:** Department of Cell Biology, Emory University, 615 Michael St, Atlanta, GA, USA, 30322; Department of Biochemistry & Biophysics, 307 Linus Pauling Science Center, Oregon State University, Corvallis, OR 97331; Linus Pauling Institute, 307 Linus Pauling Science Center, Oregon State University, Corvallis, OR 97331; School of Biological Sciences, Illinois State University, 251 S. School Street, Normal IL 61761; Department of Biochemistry, Emory University, 1510 Clifton Rd, Atlanta, Georgia, USA, 30322; Department of Neurology, Emory University, 12 Executive Park Dr NE, Atlanta, Georgia, USA, 30322

## Abstract

Impairments to the complex machinery regulating copper homeostasis lead to neurodevelopmental diseases, demonstrating the importance of copper for neuronal health and maintenance. The exact mechanisms by which the brain responds to copper deficiency following disruptions to the copper transporters ATP7A and CTR1 in conditions such as Menkes disease remain unclear, though failure to supply complex IV of the respiratory chain with copper is suspected to account for substantial pathology. Here, we studied mechanisms of copper deficiency using systems biology approaches to contrast isogenic CTR1- and COX17-deficient cells, which model copper deficiency at the level of the whole cell or complex IV, respectively. Multiomics approaches revealed distinct signaling mechanisms elicited by compartment-specific genetic defects of copper homeostasis, spanning multiple organelles and biological functions. Specifically, COX17 KO cells exhibited elevated AMPK activity and blunted mTOR activity relative to CTR1-null cells. Manipulating mTOR activity elicited inverse effects on survival in CTR1-deficient cells and flies as compared to their COX17-deficient counterparts. Increased mTOR activity and downstream protein synthesis is adaptive in models of copper deficiency but deleterious in COX17-deficient cells and flies. We propose that mTOR activation represents a resilience mechanism that fails following sustained copper deficiency and impairments to mitochondrial respiration.

**Significance:**

- Comparative proteomics reveals distinct molecular mechanisms downstream of compartmentalized copper deficiency
- COX17- and CTR1-deficient cells exhibit distinct patterns of AMPK/mTOR pathway activity
- mTOR and downstream S6K activity is protective in cellular copper deficiency but not in compartmentalized mitochondrial copper deficiency

## Introduction

Copper is a critical micronutrient with a complex system of transporters and chaperones devoted to maintaining proper copper homeostasis (Lane *et al*., 2025a). One such cuproprotein is COX17, a chaperone that donates copper to the assembly factors SCO1, SCO2, and COX11 of complex IV of the respiratory chain (Glerum *et al*., 1996; Punter and Glerum, 2003; Horng *et al*., 2004). COX17 KO is lethal in mice early in embryonic development, concomitant to gastrulation (Takahashi *et al*., 2002). Death in utero occurs at a similar timepoint following loss of the copper transporter 1 (CTR1) (Kuo *et al*., 2001; Lee *et al*., 2001), which imports copper into the cell. Knockout of the sex-linked copper transporter *ATP7A* is also embryonic lethal in males (Wang *et al*., 2012), and around half of the *mottled* mice strains (containing mutations in the locus encoding *ATP7A*) characterized to date are also embryonic lethal (Lenartowicz *et al*., 2015), with symptom severity corresponding to the impact of the mutation on copper transport and protein localization (Kim and Petris, 2007). This demonstrates the importance of functional copper homeostasis for development and highlights the specific requirement of proper copper delivery to complex IV.

Accordingly, mutations in *ATP7A* and *SLC31A1* (the gene encoding CTR1) are associated with rare childhood neurodegenerative diseases of copper deficiency (OMIM: 309400, 620306). Menkes disease, caused by defects in *ATP7A*, presents with seizures, neurodegeneration, and growth defects in the first months of life (Menkes *et al*., 1962; Tümer and Møller, 2010; Kaler, 2011). Neurological symptoms are not apparent at birth in either mouse models (Donsante *et al*., 2011; Lenartowicz *et al*., 2015; Guthrie *et al*., 2020; Yuan *et al*., 2022) or patients with Menkes disease (Kaler, 2011; Fujisawa *et al*., 2022), and copper-histidine administration is only effective if initiated in the first week or month of life in mice and humans, respectively (Fujii *et al*., 1990; Sarkar *et al*., 1993; Kaler *et al*., 2008). CTR1 deficiency presents with similar symptoms but a faster pathological timeline (Batzios *et al*., 2022; Dame *et al*., 2022; Juliá-Palacios *et al*., 2025). While mutations in SCO1/2 and COX11 are associated with diseases of mitochondrial complex IV deficiency and ATP7A/CTR1 with the diseases described above (Human Phenotype Ontology Database, https://hpo.jax.org/, Gargano *et al*., 2024), no disease or phenotype annotations exist for COX17. Of these cuproproteins, COX17 alone is categorized as a common essential gene in a larger pan-cancer screen (Arafeh *et al*., 2025; DepMap Broad, 2025), further demonstrating its obligate upstream role in complex IV function and assembly.

The exact mechanisms by which the brain responds to copper deficiency are unknown. The lack of neurological symptoms at birth and the short therapeutic window for copper delivery in Menkes disease suggests that the brain engages resilience mechanisms during this early period of neurodevelopment in response to copper deficiency that delay and/or contribute to neurological phenotypes (Lane *et al*., 2025a). We posit that this period of resilience early in life comprises compensatory responses that fail following sustained copper deficiency in the brain and subjection to new challenges induced by the transition in energy production primarily by glycolysis to mitochondrial respiration (Goyal *et al*., 2014; Kuzawa *et al*., 2014; Steiner, 2020; Oyarzábal *et al*., 2021) and concomitant increased neuronal copper demand after birth (Hatori *et al*., 2016; Chakraborty *et al*., 2022).

Failure to properly assemble complex IV and subsequent impaired mitochondrial respiration has long been hypothesized to be upstream of disease pathology in Menkes disease and patients with *SLC31A1* (CTR1) mutations (Kaler, 2013; Garza *et al*., 2022). Thus, we reasoned that comparison of cells with compartmentalized copper deficiency and differing levels of mitochondrial respiration would allow us to test predictions in vitro about mechanisms engaged in early and late stages of diseases associated with copper deficiency. We simultaneously generated isogenic KO cell lines deficient in COX17 and CTR1 (the latter described previously in Lane *et al*. (2025b)). While COX17 KO cells are unable to respire, CTR1 KO cells exhibit a partial reduction in mitochondrial respiration (Lane *et al*., 2025b), thus providing a system to model mild to severe mitochondrial phenotypes.

We previously reported that cellular and mouse models of copper deficiency exhibit increased mTOR activity and protein synthesis (Lane *et al*., 2025b), similar to reports of increased mTOR activation in diseases associated with defective mitochondrial respiration such as Leigh syndrome (Johnson *et al*., 2013; Khan *et al*., 2017). Manipulations of the mTOR pathway in these contexts introduces an enigma: while mTOR inhibition is protective in Leigh syndrome (Johnson *et al*., 2013) and cells deficient in Coenzyme-Q (Wang and Hekimi, 2021), we demonstrated that mTOR activation was adaptive in supporting the survival of copper-deficient CTR1 KO cells and rescued dendritic arborization and mitochondrial localization phenotypes in copper-deficient neurons (Lane *et al*., 2025b). We hypothesized that mTOR activation is a component of an early adaptive response to copper deficiency that ultimately fails as copper deficiency disease pathology and impaired complex IV activity progresses. We aimed to explore the role of mTOR activity in the context of moderate and extreme complex IV impairments induced by compartmentalized copper deficiency in CTR1 and COX17 KO cells, respectively, using systems biology approaches. We tested whether mTOR activity was adaptive in both contexts using pharmacogenomic approaches in CTR1 and COX17 KO cells as well as genetic epistasis experiments in *Drosophila*. We found that increased mTOR activity and downstream protein synthesis is adaptive in models of cellular copper deficiency but deleterious in COX17-deficient cells and animals. We propose that resilience to copper-deficiency cell death is tuned by a balance of protein synthesis and AMP-dependent signaling mechanisms under the control of the respiratory chain.

## Results

### Metabolic and metal phenotypes in COX17-deficient cells

We aimed to identify shared and unique pathways and signaling mechanisms in cells with compartmentalized vs. whole cell defects in copper content. We engineered isogenic SH-SY5Y cells deficient in either *COX17* (COX17 KO, lacking a copper metallochaperone essential for assembly of Complex IV/COX) or *SLC31A1* (CTR1 KO, with impaired cellular copper import and copper deficiency) (Figure 1A, 1B) by CRISPR genome editing. Wild type and COX17 KO clones were isolated simultaneously, and we confirmed that COX17 expression was abolished in the mutant clones by immunoblot (Figure 1C). To assess Complex IV activity, clones were further characterized by their survival in galactose media, preventing glycolysis, relative to glucose media. COX17 KO cells had approximately 30% survival in galactose media as compared to wild-type cells (Figure 1D). The clones with the lowest survival in galactose media were selected for further experiments. CTR1 KO cells were previously reported and described in Lane *et al*. (2025b).

**Figure 1.**
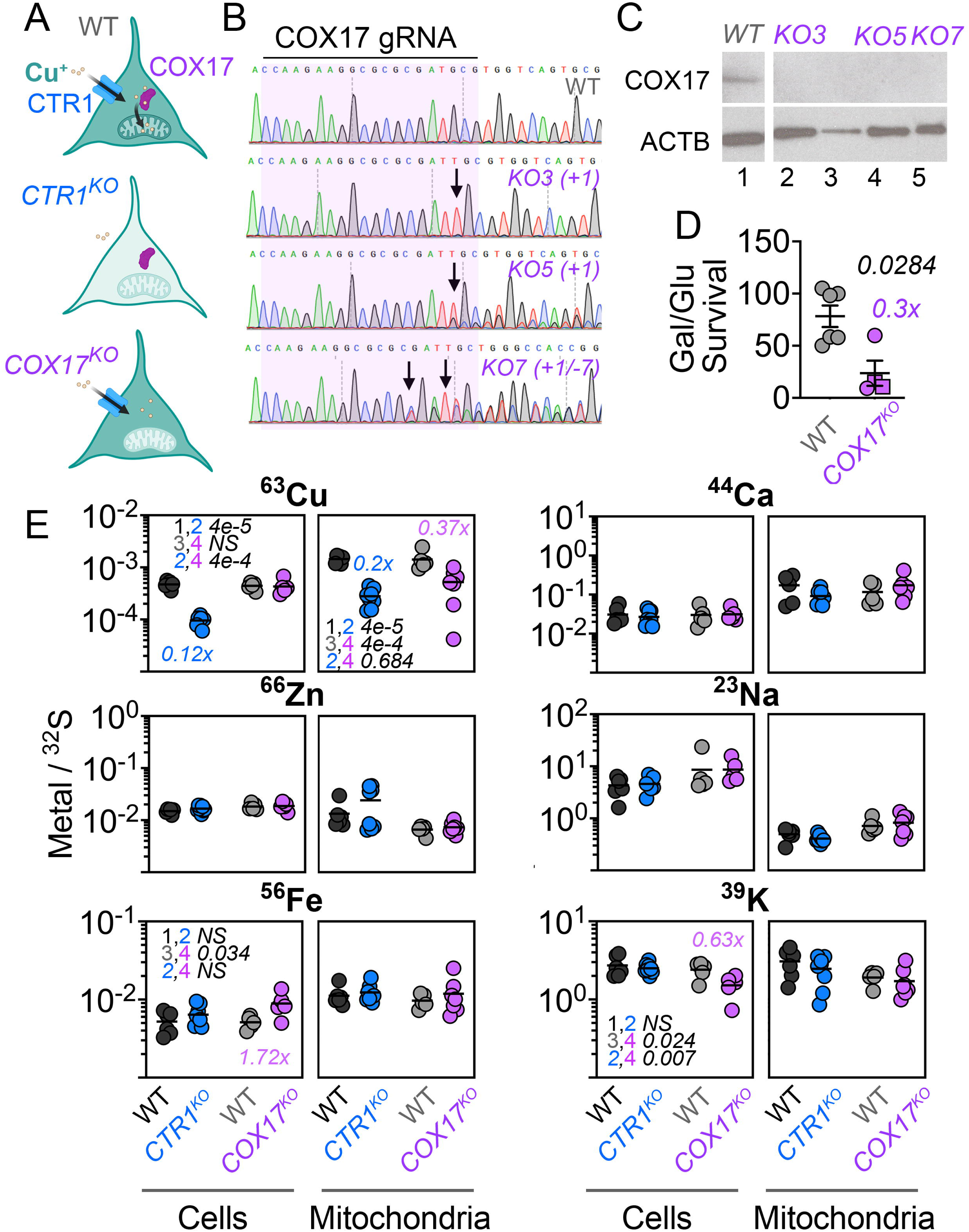
Generation and characterization of COX17-null mutant neuroblastoma cells. **A.** Diagram of compartmentalized copper deficiency in isogenic wild type, CTR1, and COX17 KO SH-SY5Y cells. Green color represents predicted copper content in cells and mitochondria. CTR1 and COX17 proteins are depicted in blue and purple, respectively. **B.** DNA sequence chromatograms of clonal isolates of wild type and three *COX17* CRISPR-edited SH-SY5Y lines. Purple boxes mark the sequence targeted by the gRNA and arrows show the mutated site. **C**. Immunoblot of cellular extracts from wild type (lane 1) and four independent *COX17*Δ*/*Δ mutant (COX17 KO, lanes 2-5) clonal cell lines probed for COX17 with beta-actin as a loading control. The unlabeled lane was for a clone that was not utilized in any other experiments. **D.** Cell survival of wild type and COX17 cell lines in normal and galactose media expressed as ratio (average ± SEM). Data are expressed as normalized to the survival of one wild type clone from each experiment. Purple text (0.3x) indicates that cell survival of COX17 cells is 1/3 of the wild-type cells in media with galactose. Black italicized numbers represent p values (unpaired mean difference two-sided permutation t-test). **E.** Metal quantification of 63Cu, 66Zn, 56Fe, 44Ca, 23Na, and 39K in whole cells or mitochondria enriched fractions in isogenic wild type and two clonal lines for both COX17 and CTR1 KO cells, normalized to 32S (average, n = 5 for wild type and n = 7-8 for COX17 and CTR1 KO cells respectively). Black italicized numbers represent p values (two-way ANOVA, followed by uncorrected Fisher’s LSD). Colored values represent relative metal levels of each mutant normalized to the corresponding wild type. There are no significant differences by treatment or genotype for 66Zn, 44Ca, or 23Na. CTR1 KO cell data in panel **D** was obtained concurrently and was reported in Lane *et al*. (2025b).

To compare the metal status of COX17- and CTR1-deficient cells, we performed ICP-MS and quantified levels of copper, zinc, iron, calcium, sodium, and potassium normalized to sulfur as described in Lane *et al*. (2025b) in both whole cells and mitochondrial-enriched fractions (Figure 1E). Total copper levels were similar in COX17 KO cells and wild-type cells, while CTR1 KO cells had a dramatic reduction in total copper levels, as previously reported (Figure 1E, 0.12x wild-type levels, Lane *et al*. (2025b)). Both mutants exhibited similar reductions in mitochondrial copper between 2-to 2.7-fold compared to wild-type cells (Figure 1E; compare columns 2 and 4). COX17 KO cells exhibited no changes in copper transporter ATP7A or the copper chaperone CCS by immunoblot but had decreased protein abundance of CTR1 compared to wild-type cells (Figure 1S). COX17-deficient cells also exhibited a 1.7-fold increase in total iron levels (Figure 1E; compare columns 3 and 4) and 1.5-fold decrease in total potassium levels with no change in sodium levels (Figure 1E; compare columns 3 and 4). No other metals were altered in CTR1 KO cells. These results show that COX17 and CTR1 KO cells have compartmentalized alterations of copper content.

The similar reduction of copper content in mitochondrial fractions prompted us to characterize the abundance, assembly, and function of the electron transport chain. We previously reported that CTR1 KO cells have reduced levels of Complex IV (see blue bar in Figure 2C) and impaired assembly of Complexes I, III, and IV (Lane *et al*., 2022). As both CTR1 and COX17 KO cells exhibit reduced mitochondrial copper, and COX17 is essential for the assembly of Complex IV, we predicted these same phenotypes would be present in COX17-deficient cells. Indeed, COX17 KO cells had decreased levels of Complex IV (0.26x wild-type levels), which is half of the complex IV content in CTR1 KO cells (0.56x wild-type levels). These changes in protein levels were accompanied by altered migration of Complexes I, III, and IV in blue native electrophoresis (Figure 2A-C). Interestingly, COX17 KO cells also exhibited reduced Complex I as compared to wild-type cells (0.48-fold, Figure 2A-C), which was not observed in CTR1 KO cells (see blue bar in Figure 2C, Lane *et al*. (2025b)). We functionally confirmed these biochemical phenotypes measuring respiratory chain function. We used two approaches: the Resipher system, which utilizes platinum organo-metallic oxygen sensors to measure whole-cell mitochondrial respiration (as defined by its abrogation with a mix of rotenone plus antimycin) over the course of several days while maintaining cells in their own milieu (Grist *et al*., 2010; Wit *et al*., 2023), and Seahorse, which measures oxygen consumption rate (OCR) and extracellular acidification rate (ECAR) over a few hours in serum-free media and reports multiple parameters of mitochondrial respiration. By either Resipher or Seahorse, there is no detectable mitochondrial respiration in COX17-deficient cells (Figure 2D-F, purple markers). This is in contrast to CTR1 KO cells, which have approximately 35% OCR levels as compared to wild type as shown by Resipher (Figure 2D, blue markers), reproducing our previously reported findings (Lane *et al*. (2025b); blue box plots in Figure 2F included for reference). Unsurprisingly given their lack of mitochondrial respiration, COX17 KO cells have increased glycolysis and glycolytic capacity and reduced glycolytic reserve as compared to wild-type cells (Figure 2G-H, purple markers), similar to our previously reported results in CTR1 KO cells (see blue bars, Figure 2H). Altogether, this indicates that while cellular copper deficiency is unique to CTR1 KO cells, CTR1 and COX17 deficiency produce similar mitochondrial copper deficits and elicit increased glycolysis and impaired respiration (Figure 2I), a phenotype that is more exaggerated in COX17 KO cells.

**Figure 2.**
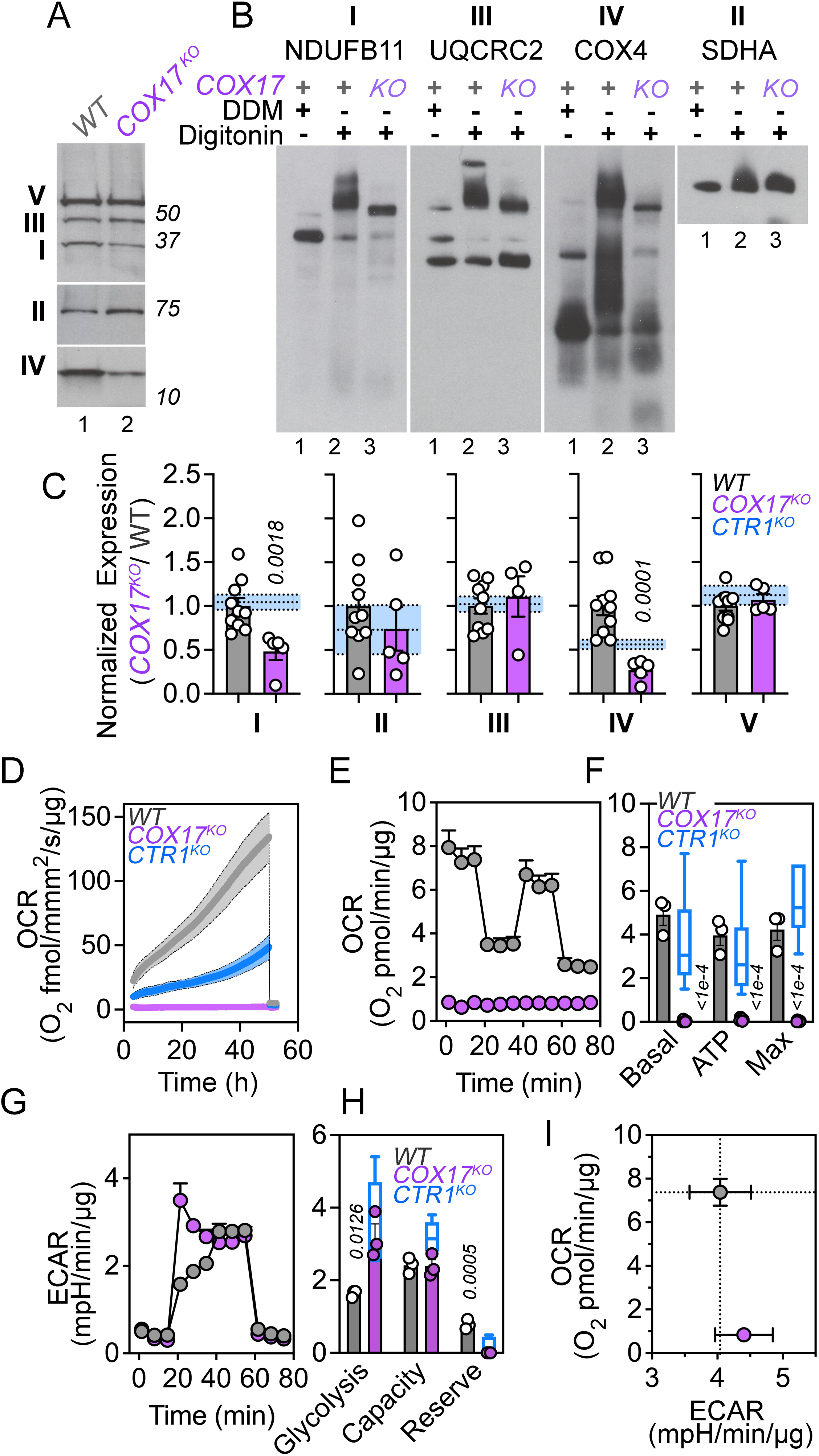
COX17 KO cells exhibit impaired electron transport chain assembly and respiration and increased glycolysis. **A-C**. Respiratory chain protein levels in mitochondrial fractions from wild type and COX17 KO cells probed by immunoblot with OxPhos antibody mix (**A**) or blue native electrophoresis and immunoblot probed with antibodies against Complex, I, II, III, and IV (**B**). For native gel immunoblots, samples were solubilized in either DDM or digitonin to dissolve or preserve supercomplexes, respectively (Wittig *et al*., 2006; Timón-Gómez *et al*., 2020). Complex II was used as a loading control as it does not form respiratory supercomplexes (Iverson *et al*., 2023). **C**. Immunoblot quantification from **A** showing normalized protein abundance to wild-type cells (average ± SEM; each dot is an independent biological replicate). Italicized numbers represent p values analyzed by unpaired mean difference two-sided permutation t-test. Blue bars represent the average ± SEM of previously reported CTR1 KO cell data that was obtained concurrently (Lane *et al*., 2025b). **D**. Resipher respiration rates in wild type, COX17 KO, and CTR1 KO cells over 48 hours, followed by termination of the assay by the addition of rotenone plus antimycin (R+A). average ± SD, n = 12 technical replicates. **E-I.** Metabolic activity in wild type and COX17 KO cells measured by the Seahorse Mito Stress Test (**E-F**, n = 3 of each genotype) and Glycolysis Stress Test (**G-H**, n = 3 for wild type and two COX17 KO clonal cell lines). Data are presented normalized to protein (average ± SEM) and analyzed by unpaired mean difference two-sided permutation t-test (italicized numbers represent p values). **I**. Metabolic map of basal OCR and ECAR from the Mito Stress Test. All data are presented as average ± SEM. Blue box-and-Tukey whiskers plot in F and H represent of previously reported CTR1 KO cell data that was obtained concurrently (Lane *et al*., 2025b).

### Comparative proteomics reveals distinct molecular mechanisms downstream of COX17 and CTR1 deficiency

To identify molecular pathways and signaling mechanisms responsive to COX17 and CTR1 deficiency, we used quantitative mass spectrometry of total and phosphorylated proteomes of whole cell extracts by Tandem Mass Tagging (TMT) of COX17 and CTR1 KO cells simultaneously (Figure 3, Figure S31, Supplemental Data File 1). In total, we quantified 8,986 proteins and 19,082 phosphopeptides (from 4066 proteins) across wild-type cells, two CTR1 KO clones (Lane *et al*., 2025b), and two COX17 KO clones (Figure 3A, volcano plots of COX17 KO vs wild-type cells; Figure 3S1, Venn diagram summary). The integrated proteomes, consisting of proteins and phosphopeptides with differential abundance in each genotype (q < 0.05 and fold of change ≥ 1.5, one-way ANOVA followed by Benjamini-Hochberg FDR correction), segregated by genotype, with COX17 KO cells clustering the greatest Euclidean distance away from wild type and CTR1 KO clones (Figure 3B, PCA). The CTR1 KO proteome and phosphoproteome were analyzed and key targets validated by immunoblot in Lane *et al*. (2025b). Overall, we identifed 838 proteins and 1243 phosphoresidues (corresponding to 763 unique proteins) that exhibited differential abundance in COX17 KO cells compared to wild-type cells, in marked contrast to the 210 proteins and 224 phosphoresidues (corresponding to 161 unique proteins) with differential abundance in CTR1 KO cells compared to wild-type cells (Figure 3S1).

**Figure 3.**
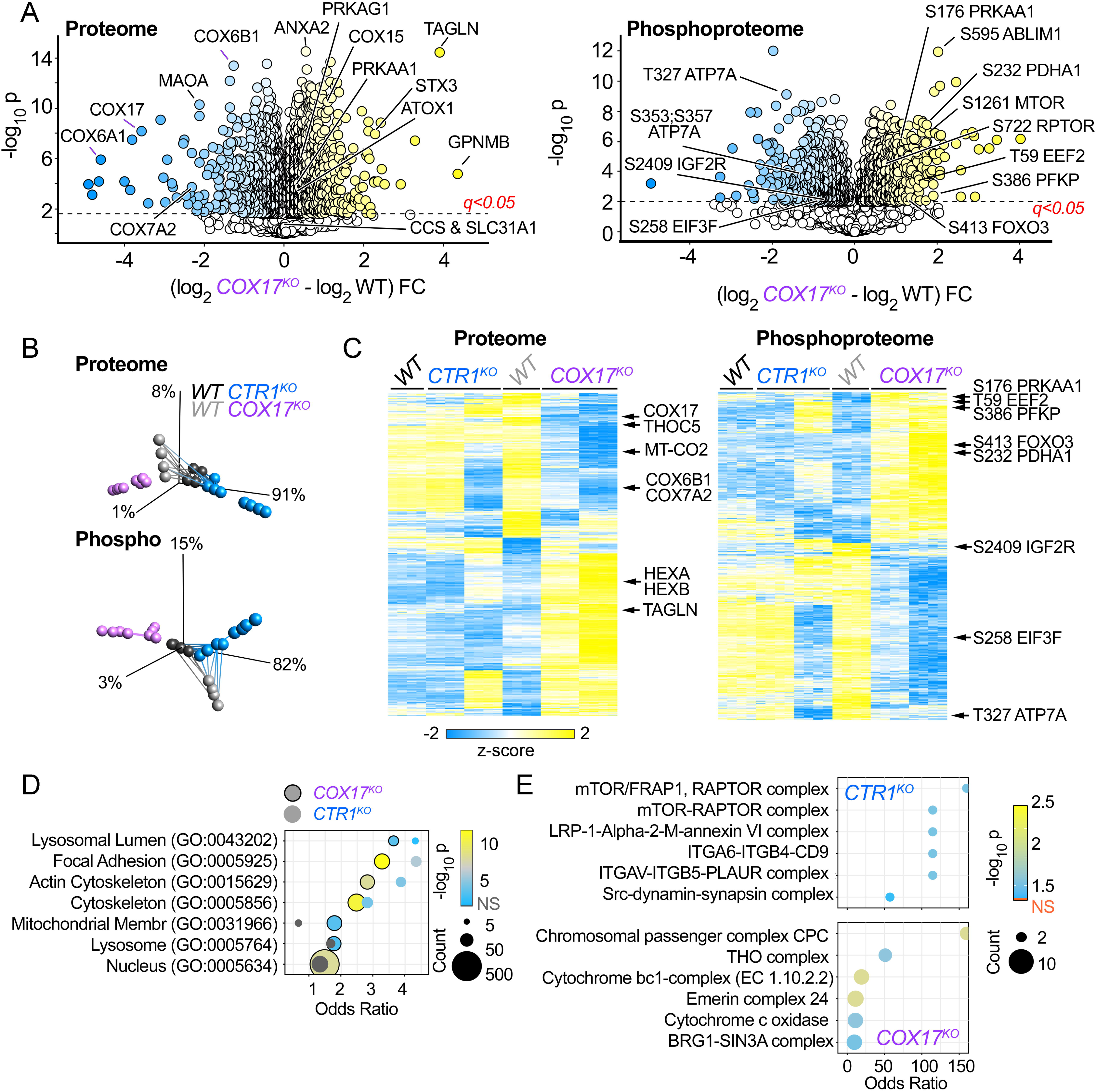
Ontological and signaling differences inferred from isogenic COX17-and CTR1-null proteomes and phosphoproteomes. **A.** Volcano plots of the COX17 KO cell proteome and phosphoproteome, where yellow dots represent proteins or phosphoproteins whose level is increased in KO cells and blue dots represent decreased expression in KO cells (n = 4 for wild-type cells and n = 4 for KO cells in two independent clones, KO3 and KO5). **B.** Principal component analysis (PCA) of the integrated proteome (consisting of the proteome and phosphoproteome) from two wild type isolates, two CTR1 KO, and two COX17 KOs clonal lines shown in **A**. PCA was performed using all proteome or phosphoproteome hits where differential expression is significant with q < 0.05 and fold of change ≥ 1.5 (one-way ANOVA followed by Benjamini-Hochberg FDR correction). Data clustering was performed by Euclidean distance (lines between symbols). TMT labeling and mass spectrometry of all groups was performed concurrently, and CTR1 KO cell data was previously reported in Lane *et al*. (2025b). **C.** Hierarchical clustering of the whole proteome and phosphoproteome from the cells described in **B**. Proteins or phosphopeptides were selected for hierarchical clustering analysis by comparing the wild type and COX17 proteomes and phosphoproteomes and thresholding by q < 0.05 and fold of change ≥ 1.5 (t-test followed by Benjamini-Hochberg FDR correction). Note differences between the COX17 KO-driven proteome and phosphoproteome with the parallel CTR1 KO dataset. **D-E.** Gene ontology (GO) analysis of merged datasets of differentially expressed proteins and protein phosphopeptides (q < 0.05 and FC ≥ 1.5) for either CTR1 or COX17 mutants, presented by bubble plot. GO Cellular Component (CC) terms for the COX17 KO integrated proteome dataset were selected from the top 25 terms with the lowest corrected p value (Ashburner *et al*., 2000; Gene Ontology Consortium *et al*., 2023); see Supplemental Data File 1 for the full list of terms). The corresponding CC term values for CTR1 KO cells were plotted alongside the COX17 values, irrespective of their p value ranking. COX17 markers are outlined in black, and CTR1 markers have no outline. Dot size indicates the number of genes annotated to each term and color indicates −log10 p value. **E.** Bubble plots showing independently top ranked CORUM annotated protein complexes for each mutant dataset, selected by lowest corrected p value. Fisher exact test followed by Benjamini-Hochberg correction. CTR1 KO cell data in panels **B-E** was obtained concurrently and was reported in Lane *et al*. (2025b).

To explore the pathways and ontologies associated with proteins and phosphopeptides with differential abundance in the combined proteome and phosphoproteome of COX17 KO cells (1458 proteins) and CTR1 KO cells (345 proteins), defined as the integrated proteome, we queried the NCAST BioPlanet discovery resource (Huang *et al*., 2019), Gene Ontology (GO) Cellular Component annotations (Ashburner *et al*., 2000; Gene Ontology Consortium *et al*., 2023), and the CORUM database (Tsitsiridis *et al*., 2023) with the ENRICHR tool (Chen *et al*., 2013; Kuleshov *et al*., 2016; Xie *et al*., 2021). The top enriched terms identified from the BioPlanet resource for each genotype were non-overlapping with the exception of BDNF signaling (Figure 3S2, Supplemental Data File 1), demonstrating distinct proteome phenotypes between COX17- and CTR1-deficient cells.

We next performed gene ontology (GO) analysis of the COX17 KO integrated proteome (Figure 3D, Supplemental Data File 1). The top GO Cellular Component (CC) terms included ontologies associated with the lysosome (Lysosomal Lumen GO:0043202, q = 2.76e-4; Lysosome GO:0005764, q = 7.52e-4), cytoskeleton and actin (Focal Adhesion GO:0005925, q = 1.26e-13; Actin Cytoskeleton GO:0015629, q = 1.68e-9; Cytoskeleton GO:0005856, q = 1.48e-12), the Mitochondrial Membrane (GO:0031966, q = 3.29e-4), and the Nucleus (GO:0005634, q = 3.35e-11). We searched for these same ontologies in the CTR1 KO integrated proteome and observed that the representation of these ontologies in CTR1 KO cells was less prominent as defined by the number of annotated proteins and the q value per ontology (Figure 3D, compare sizes). In fact, of these ontologies, only the Lysosomal Lumen (q = 4.3e-2), Focal Adhesion (q = 4.58e-7), Actin Cytoskeleton (q = 2.68e-5), and Cytoskeleton (q = 6.5e-5) terms were also significantly enriched in CTR1 KO cells (Figure 3D, Supplemental Data File 1).

Ten times more proteins annotated to the Mitochondrial Membrane were identified in the COX17 KO integrated proteome as compared to the CTR1 KO proteome (72 vs. 7, Figure 3D, Supplemental Data File 1). This included decreased abundance of multiple subunits of Complex IV (COX) as well as COX17 itself. Levels of these Complex IV subunits (COX5B, COX6A1, COX6B1, COX6C, COX7A2, COX7B) were also reduced but to a lesser degree in CTR1 KO cells. COX17-deficient cells also exhibited increased phosphorylation of PFKP, an isoform of phosphofructokinase-1 and a rate-limiting enzyme in glycolysis (Yi *et al*., 2019), at S386 and at multiples sites of PDHA1, a component of the pyruvate dehydrogenase complex (pT131, S232, pS293/S300; Sugden and Holness (2003); Figure 3C). CTR1-deficient cells have reduced phosphorylation of PFKP at this same site, which has been shown to be phosphorylated by AKT and ultimately prevents its degradation and promotes glycolysis (Zhou *et al*., 2025). Phosphorylation of PDHA1 at S232, S293, or S300 by pyruvate dehydrogenase kinase 1 (PDK1) inactivates the pyruvate dehydrogenase complex and prevents the conversion of pyruvate into acetyl-CoA, slowing the Krebs cycle (Roche *et al*., 2001). Both COX17- and CTR1-null cells had 1.5x wild-type levels of PDHA1, but no changes in phosphorylation were observed in CTR1-deficient cells. These changes in phosphorylation of PFKP and PDHA1 and reduced abundance of COX subunits in COX17-deficient cells and the more moderate changes in CTR1-deficient cells are consistent with the respiratory defects and increase in glycolysis we identified in these mutants (Figure 2). Interestingly, COX17 KO cells had increased phosphorylation of the copper transporter ATP7A at T327, a site which is only phosphorylated in the presence of cytoplasmic copper (Veldhuis *et al*., 2009) and for which mutations increase cell-autonomous copper accumulation and retention (Fieten *et al*., 2016). No changes in ATP7A phosphorylation were observed in CTR1 KO cells. While we observed an increase in STEAP3 of 2.2-fold in COX17 KO cells by TMT-MS, consistent with the increased iron levels in these cells (Figure 1E), we did not reproduce this by immunoblot (Figure 4A-C). Overall, the proteome and phosphoproteome corroborate the disrupted bioenergetic balance and metallobiology we observed in COX17 deficiency and demonstrate unique molecular and signaling mechanisms engaged by deficiency of COX17 or CTR1.

**Figure 4.**
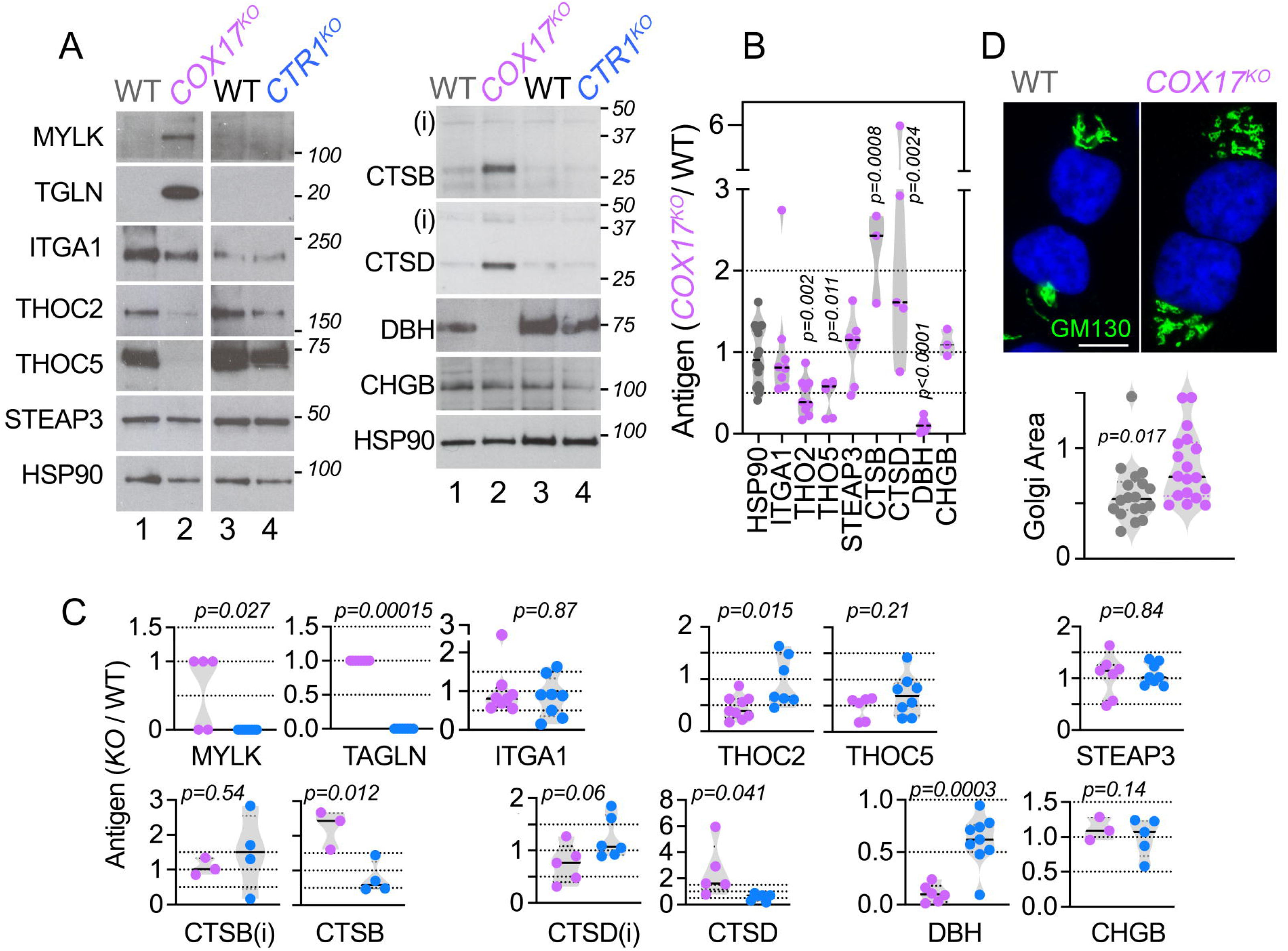
Immunoblots of GO Term-selected proteins show divergent phenotypes in isogenic COX17- and CTR1-null cells. **A-C**. Immunoblots of whole cell extracts from wild type (gray), two COX17 KO clonal lines (purple), and two CTR1 KO clonal lines (blue) probed for proteins involved in actin filaments or the cytoskeleton (MYLK, TALGN, ITGA1), components of the THO complex (THOC2, THOC5), the metalloreductase STEAP3, and secretory proteins (CTSB, CTSD, CHGB, and DBH; immature cathepsins are indicated with i). HSP90 was used as a loading control. Immunoblots were quantified by normalizing protein abundance to wild-type cells. The left column of images are from the same immunoblot and cropped for clarity. **B**. Quantification of COX17 KO protein abundance relative to wild type. Italicized numbers represent p values analyzed by unpaired mean difference two-sided permutation t-test. **C.** Quantification of COX17 and CTR1 KO protein abundance relative to wild-type cells. Italicized numbers represent p values analyzed by unpaired t-test with the following exceptions: MYLK and TGLN were analyzed by Fisher Exact Probability Test and CHGB was analyzed by Mann Whitney test. **D**. Immunofluorescence staining of WT and COX17 KO cells stained with the Golgi complex marker GM130 (green) and DAPI (blue) and quantification of Golgi area (pixels, each dot represents one cell). p value analyzed by unpaired t-test.

### Divergent phenotypes across multiple ontologies and organelles in the proteome and metabolic transcripts of COX17- and CTR1-deficient cells

As several of the top terms identified by GO analysis in both COX17 and CTR1 KO cells were related to the cytoskeleton and actin, we further probed these pathways by immunoblot. COX17-deficient cells exhibited increased abundance of the actin cross-linking protein and tumor suppressor transgelin (TAGLN, Shapland *et al*. (1993); Jimenez Jimenez *et al*. (2025), 15-fold increase) and the actin-binding protein and tumor suppressor myosin light chain kinase (MYLK, Kohama *et al*. (1996), 4-fold increase), both of which were 0.5x wild-type levels in CTR1 KO cells (Figure 3D, Figure 3S1, Supplemental Data File 1, q = 5.4e-7 and 8.8e-5, respectively). TAGLN can be induced by TGF-β (Chen *et al*., 2019) and p53 (Tsui *et al*., 2019). This increased abundance of TAGLN in COX17 KO cells is consistent with the upregulation of TGF-B1-induced transcript 1 protein (TGFB1I1) in COX17 KO (2.2-fold) but not CTR1 KO cells (Supplemental Data File 1). Abundance of TP53I3, tumor protein P53 inducible protein 3, is also increased 4.4-fold in COX17-null cells but is decreased by 1.8-fold in CTR1-deficient cells; TP53I3 phosphorylation at S809 and S862 is reduced by approximately half in COX17 cells compared to wild type (Supplemental Data File 1). We also observed a notable difference in the adherence of these mutants in culture, where COX17 KO cells were markedly more adherent and CTR1 KO cells far less adherent than wild-type cells. Additionally, four proteins differed in both abundance and phosphorylation in COX17 and CTR1 mutants (Figure S31); two of these proteins, ABLIM and TRIO, bind to and/or regulate the actin cytoskeleton (Roof *et al*., 1997; Seipel *et al*., 1999). Both proteins exhibit opposite patterns of abundance in COX17 and CTR1 mutant cells: ABLIM and TRIO abundance and phosphorylation at multiple sites are increased in COX17-null cells and decreased in CTR1-null cells (Figure S31). Altogether, this suggests that while actin and cytoskeleton-related pathways are altered in both COX17- and CTR1-null cells with partial overlap, distinct molecular mechanisms are engaged in response to each mutation.

We also explored proteins with differential abundance annotated to the lysosome. Lysosomal proteins with increased abundance in COX17 KO cells compared to wild type include multiple cathepsins (CTSA, CTSB, and CTSD; 1.6 to 2.0-fold increase) and both subunits of beta-hexosaminidase A (HEXA and HEXB, 1.6-fold increase); these proteins had similar abundance in CTR1 KO and wild-type cells (Figure 3C, Figure 3S1, Supplemental Data File 1). HEXA and HEXB are hexosaminidases involved in the processing of glycosphingolipids (Liu *et al*., 2018; Quinville *et al*., 2021). Cathepsins are a family of proteases with a variety of functions important for cellular homeostasis, immune responses, autophagy, and development (Yadati *et al*., 2020). We confirmed increased levels of mature CTSB and CTSD in COX17 KO but not CTR1 KO cells by immunoblot (Figure 4A-C). We observed no changes in the immature cathepsins (Figure 4A-C). We also measured the levels of the secretory enzyme dopamine beta-hydroxylase (DBH). DBH converts dopamine to norepinephrine in a copper-dependent manner, and copper levels have also been reported to affect DBH secretion (Schmidt *et al*., 2018). Both COX17 and CTR1 mutants exhibit reduced abundance of DBH by immunoblot, but COX17 KO cells have significantly less DBH than CTR1 KO despite having normal total copper levels (Figure 4A-C, Supplemental Data File 1). Importantly, MYLK or MLCK has an established role in exocytosis and secretion. MYLK, which is increased in COX17 KO cells and decreased in CTR1 KO cells, has been shown to promote the release of catecholamines like dopamine (Kumakura *et al*., 1994). Both mutants exhibited normal levels of chromogranin B, another secretory protein, indicating that this reduction in DBH and increased levels of cathepsins are not indicative of global defects in late stages of the secretory pathway (Figure 4A-C). We also observed reduced phosphorylation of insulin-like growth factor 2 receptor/Cation-Independent Mannose-6-Phosphate Receptor (IGF2R/CI-M6PR) at S2409 in COX17 KO cells (0.5x wild-type levels; Figure 3C, Supplemental Data File 1). IGF2R/CI-M6PR is required for the sorting of lysosomal enzymes from the trans-Golgi network to late endosomes. However, the functional role of the S2409 phosphorylation has not been elucidated (Méresse and Hoflack, 1993; Ghosh *et al*., 2003). These changes in the levels of proteins that traverse the Golgi complex prompted us to assess if there were morphological alterations of this compartment. Indeed, we observed increased area occupied by the Golgi complex by immunofluorescence in COX17 KO cells as compared to wild type (Figure 4D), which we speculate may be linked to increased levels of lysosomal proteins and other secretory proteins.

To expand mechanistic discovery in our rich datasets, we queried the CORUM database (Tsitsiridis *et al*., 2023) to identify protein complexes significantly enriched in the integrated proteome and phosphoproteome of COX17- and CTR1-deficient cells. Of the top six most enriched complexes for the COX17 KO cells, two are respiratory complexes and the remaining four complexes are nuclear (chromosomal passenger complex or CPC, THO complex, emerin complex, and BRG1-SIN3A complex). Six components of Complex III (CYC1, UQCR10, UQCRQ, UQCRB, UQCRC1, and UQCRC2) increased in abundance in COX17 KO (1.5x to 1.8x wild-type levels). Consistent with our respiration measurements and immunoblots of Complex IV (Figure 2), six Complex IV subunits decreased in expression between 0.03-0.6x wild-type levels (COX6B1, COX6A1, COX7A2, COX5B, COX6C, and COX7B). All four components of the chromosomal passenger complex (CPC) (AURKB, INCENP, BIRC, CDCA8), a key regulator of mitosis (Carmena *et al*., 2012), had reduced abundance in COX17 KO cells (0.6x wild-type levels) with no change in the CTR1 KO cells. Similarly, four of the five components of the THO complex (THOC1, THOC2, THOC5, THOC7), which is involved in the formation of messenger ribonucleoparticles (Jimeno and Aguilera, 2010), were present at ∼0.6x wild-type levels in COX17 KO but were not altered in CTR1 KO cells (Figure 3S1). Phosphorylation of THOC2 at T1285, T1289, and S1393 in COX17 KO cells was approximately half the levels of wild type, and phosphorylation of INCENP was also reduced (0.3x wild-type levels). Decreased abundance of THOC2 and THOC5 in COX17 KO but not CTR1 KO cells was verified by immunoblot (Figure 4A-C). The mTOR-RAPTOR complex was enriched specifically in the CTR1 KO cells (q = 0.023, Figure 3E), consistent with our previous reports of increased mTOR-S6K activation in copper deficiency (Lane *et al*., 2025b).

As an orthogonal approach to proteomics, we performed Nanostring nCounter transcriptome analysis with the Metabolic Pathways panel to measure the expression of approximately 800 curated metabolism-annotated genes in COX17 and CTR1 KO cells (Figure 5, Supplemental Figure 5S). CTR1 KO data was previously reported in Lane *et al*. (2025b). We detected 261 differentially expressed genes in COX17 KO cells and 222 differentially expressed genes in CTR1 KO cells (q < 0.05 and fold of change ≥ 1.5, one-way ANOVA followed by Benjamini-Hochberg FDR correction, Supplemental Figure 5S). Differentially expressed curated metabolic transcripts segregated genotypes by PCA, with COX17 and CTR1 KO cells clustering away from wild-type cells (Figure 5B). We further narrowed our analysis to genes involved in bioenergetics (specifically the respiratory chain, pyruvate metabolism, and glycolysis), the AMPK and mTOR pathways, and the lysosome (Figure 5D-F). Similar changes in expression are observed in both COX17 and CTR1 KO cells (Figure 5D-F), but changes were significantly more pronounced in AMPK/mTOR and lysosomal annotated transcripts in COX17 KO cells as compared to CTR1 KO cells, as revealed by principal component 1 of the annotated transcripts in each annotation (Figure 5G, see Figure 6S). Corroborating the proteomics data, we observed increased expression of CTSA and CTSD in COX17 but not CTR1 KO cells (Figure 5F, Supplemental Figure 5S). HEXA and HEXB were upregulated modestly in CTR1-null cells (1.3-1.5x wild-type levels) and dramatically increased in COX17-deficient cells (2.1-2.2x wild-type levels, Figure 5F, Supplemental Figure 5S). Three AMPK subunits were upregulated in COX17-null cells with no change in CTR1 KO cells (PRKAA1, 1.46x wild-type levels; PRKAG1, 1.38x wild-type levels; PRKAG2, 2.14x wild-type levels (Figure 5E, Supplemental Figure 5S). Together, the integrated proteomes and curated metabolic transcripts from COX17- and CTR1-deficient cells demonstrate unique mechanisms in each genotype impacting mitochondrial proteins and respiratory complexes, actin-binding and actin regulatory proteins, multiple organelles, and the AMPK and mTOR signaling pathways.

**Figure 5.**
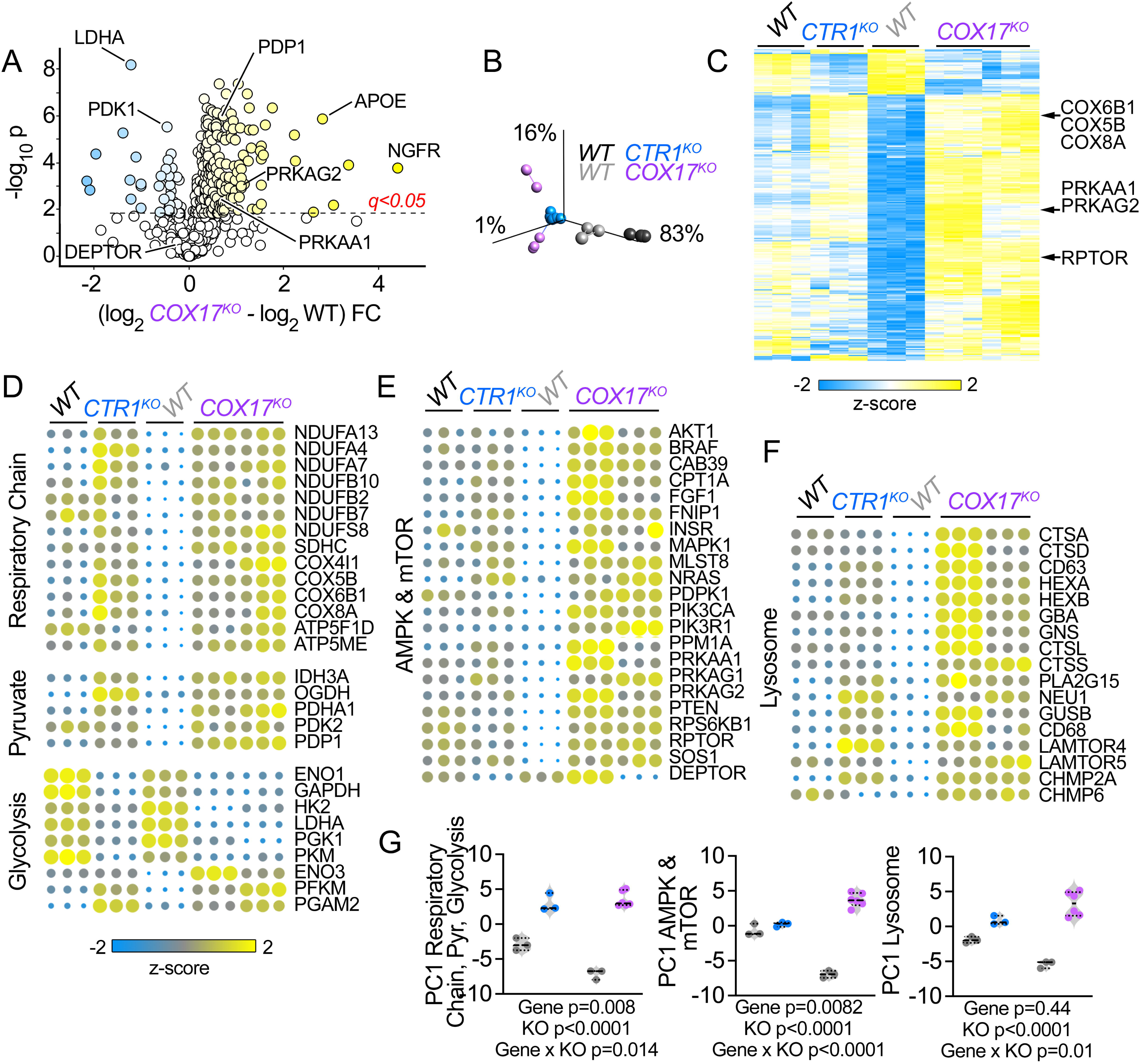
NanoString curated metabolic transcripts are differentially affected in isogenic COX17- and CTR1-null cells. **A.** Volcano plot of the 768 NanoString Metabolic Pathways transcripts from wild type (gray) or two COX17 KO (purple) clonal lines for genes that are differentially expressed at a significance of q < 0.05 (t-test followed by Benjamini-Hochberg FDR correction) and fold of change ≥ 1.5. See Supplemental Data File 1 for source data. **B.** Principal component analysis of curated metabolic transcripts from wild type (gray), COX17 KO (purple), or CTR1 KO (blue symbols) cells thresholded by q < 0.05 (one-way ANOVA followed by Benjamini-Hochberg FDR correction). Data clustering was performed by Euclidean distance (lines between symbols). **C.** Hierarchical clustering of the curated metabolic transcripts comparing wild type, CTR1 KO, and COX17 KO cells, showing differentially expressed mRNAs selected by a q < 0.05 and a fold of change ≥ 1.5 in COX17 KO cells (t-test followed by Benjamini-Hochberg FDR correction). **D-G.** Heat maps and principal component analysis of curated metabolic transcripts of genes differentially expressed in COX17 cells and annotated to the COX17 ontologies in Fig. 3D-E and S3. **G.** Depicts Principal Component 1 (PC1) for each set of genes in **D-F** (two-way ANOVA followed by Bonferroni’s multiple comparisons tests). Wild type, CTR1, and COX17 KO transcriptomes were obtained concurrently. CTR1 KO cell data was previously reported in Lane *et al*. (2025b).

**Figure 6.**
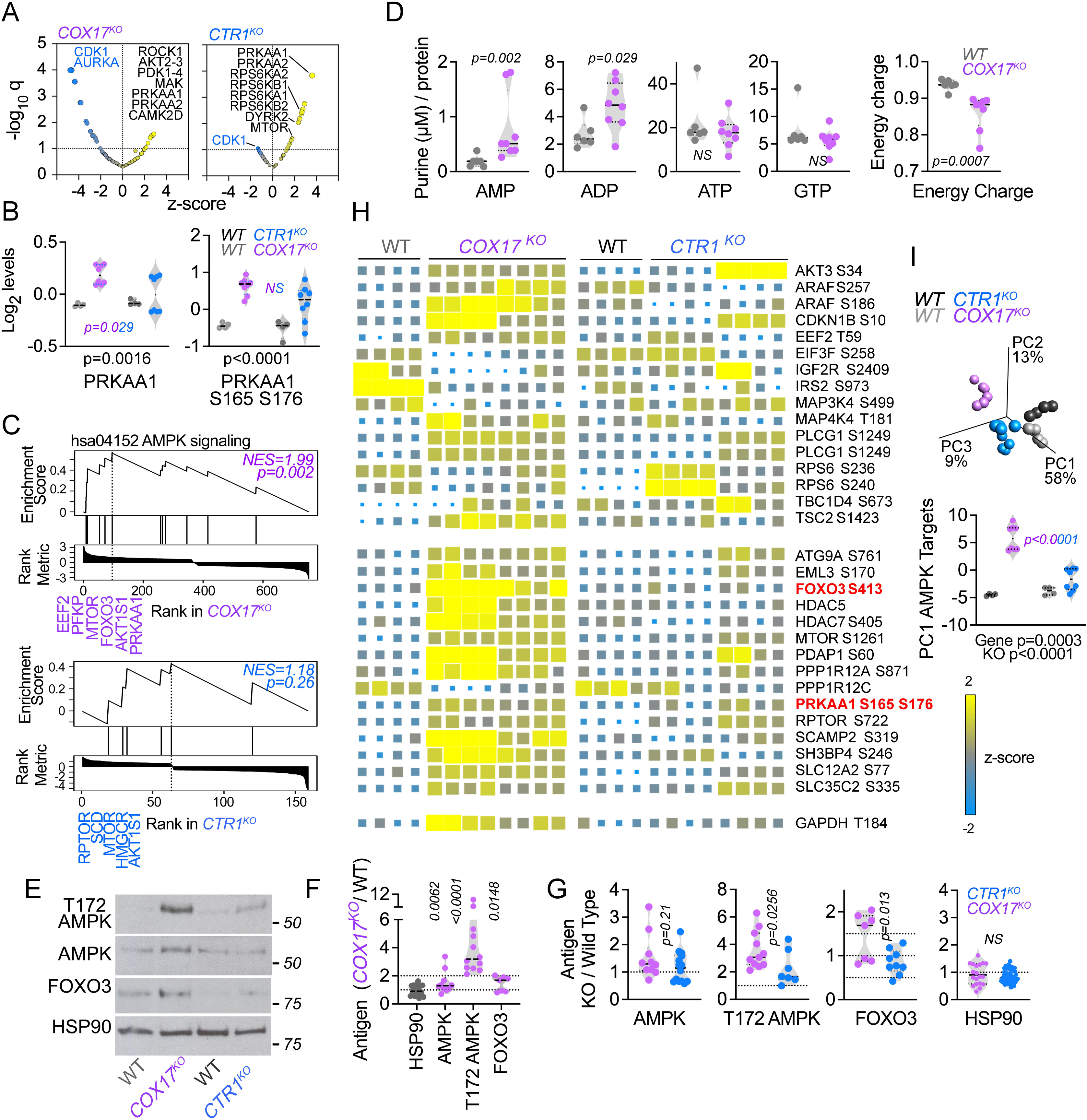
Differential AMPK pathway activity in COX17- and CTR1-null cells. **A.** Kinase activity scores generated from COX17 KO and CTR1 KO phosphoproteomes with the Kinase-Substrate Enrichment Analysis (KSEA) app (Wiredja *et al*., 2017). **B.** Mass spectrometry quantification of total and phosphorylated AMPK (PRKAA1 pS165/S176) from two CTR1 KO clonal lines (blue symbols), two COX17 KO clonal lines (purple symbols), and their corresponding wild type clonal lines (dark gray and light gray, respectively). Two-way ANOVA followed by Bonferroni’s multiple comparisons test. **C.** Gene set enrichment analysis (GSEA) of the phosphoproteomes for either COX17 or CTR1 KO cells. Graphs show normalized enrichment score (NES) for AMPK signaling (hsa04152). Note the distinct phosphopeptides with the highest NES in COX17 vs CTR1 KO cells. p value calculated with weighted Kolmogorov-Smirnov-like statistic according to Wang *et al*. (2017a). **D**. HPLC quantification of purine nucleotides from wild type and two COX17 KO clonal lines (analyzed by Mann Whitney test, n = 6-8 per genotype). Energy charge was calculated as ([ATP] + 0.5[ADP]) / ([ATP]+[ADP]+[AMP]) (Atkinson and Walton, 1967). **E-G**. Immunoblots of whole cell extracts from isogenic wild type, two COX17 KO, and two CTR1 KO clonal lines probed for total and phosphorylated AMPK (pT172), FOXO3, and HSP90 as a loading control. Immunoblots were quantified by normalizing protein abundance to wild-type cells. Italicized numbers represent unpaired two-sided permutation t-test as compared to WT (**F**) or between the two genotypes (**G**). **F**. Quantification of COX17 KO protein abundance relative to wild type. **G**. Quantification of COX17 and CTR1 KO protein abundance relative to wild-type cells. **H**. Heat map of AMPK substrates curated in Schaffer *et al*. (2015) whose differential expression is significant with q < 0.05 and fold of change ≥ 1.5 in COX17-null cells (t-test followed by Benjamini-Hochberg FDR correction). Each box represents a replicate. **I.** Depicts Principal Component analysis and PCA 1 (PC1) of all the genes in heat map shown in G comparing isogenic wild type, two COX17 KO clonal lines, and two CTR1 KO clonal lines. Two-way ANOVA followed by Bonferroni’s multiple comparisons tests.CTR1 KO cell data in panels **A-D** and **H-I** was obtained concurrently and was reported in Lane *et al*. (2025b).

### Altered balance of AMPK and mTOR pathway activity in COX17- and CTR1-deficient cells

To explore signaling mechanisms engaged in COX17 KO and CTR1 KO cells, we estimated kinase activity *in silico* with the Kinase-Substrate Enrichment Analysis (KSEA) tool (Casado *et al*., 2013; Wiredja *et al*., 2017). AMPK activity was significantly enriched in both mutants, while mTOR activity was significantly enriched only in CTR1 KO cells (Figure 6A, Supplemental Data File 1). Based on these kinase activity estimates and our observations of distinct changes in AMPK-mTOR pathway genes in COX17 and CTR1 KO cells, we chose to explore our proteomics data for gene products annotated to these pathways. We observed that COX17-deficient cells had increased AMPKα1/PRKAA1 levels as compared to wild type (1.22-fold), which was significantly higher than CTR1 KO cells (Figure 6B). Total AMPKα was increased in both mutants, with significantly higher levels in COX17-null cells as compared to CTR1, as confirmed by immunoblot (1.54x wild-type levels in COX17 KO, 1.17x wild-type levels in CTR1 KO, Figure 6E-G, 4S). Both mutants had increased AMPK α1/PRKAA1 S165/S176 phosphorylation (2.08x wild-type levels in COX17, 1.62-fold wild-type levels in CTR1) (Figure 6B, 4S). Increased AMPKα1/2 T183/T172 phosphorylation, which increases AMPK activity (Steinberg and Carling, 2019), was observed by immunoblot in both mutants but was significantly higher in COX17-deficient cells (Figure 6E-G, 4S; COX17 KO have 1.8x CTR1 KO levels). Increased AMPK abundance and phosphorylation in COX17-deficient cells is consistent with their increased levels of AMP (4.1x wild-type levels) and decreased energy charge (0.92x wild-type levels) (Figure 6D; Herzig and Shaw (2018)), similar to what we previously reported for CTR1 KO cells (Lane *et al*., 2025b).

While both mutants had increased AMPK S165/S176 phosphorylation, we observed increased phosphorylation of multiple downstream targets of AMPK in COX17-deficient cells as compared to CTR1-deficient cells. For example, phosphorylation of eukaryotic elongation factor 2 (eEF2) at T57/T59 was increased in COX17 KO cells (2.2-10.8x wild-type levels) but not in CTR1 KO cells (Figure 6H). AMPK activity is known to induce phosphorylation of eEF2 by activating eEF2K directly or upstream inhibition of mTORC1 (Johanns *et al*., 2017). COX17 KO cells also have reduced phosphorylation of EIF3F at S258 (0.54-fold, q = 0.033, Figure 6H). EIF3F is a subunit of the eukaryotic translation initiation factor 3 (eIF3) complex that interacts directly with mTOR and S6K (Holz *et al*., 2005; Harris *et al*., 2006), and its phosphorylation at S258 plays a role in translational repression specifically during M phase (An *et al*., 2020). We also observed increased phosphorylation of the transcription factor FOXO3 in COX17 KO cells. The FoxO family is known to induce stress response pathways and are important regulators of metabolism, proliferation, and cell fate (Rodriguez-Colman *et al*., 2024); FOXO3 specifically blocks activity of mTORC1 (Chen *et al*., 2010). We observed increased phosphorylation of FOXO3 at S413 in COX17 but not CTR1 KO cells (2.4x wild-type levels, Figure 6F, Figure 6S), a site known to be phosphorylated by AMPK without affecting FOXO3 nuclear localization (Greer *et al*., 2007; Wang *et al*., 2017b). COX17 KO cells also exhibited a modest increase in FOXO3 phosphorylation at S253 (1.4x wild-type levels, Supplemental Data File 1), an inhibitory site phosphorylated by Akt that prevents nuclear import and causes FOXO3 to be retained in the cytoplasm (Brunet *et al*., 1999; Plas and Thompson, 2003; Dobson *et al*., 2011). We observed a modest increase in total FOXO3 by TMT (1.2-fold) and immunoblot (Figure 6E-G).

Importantly, AMPK signaling was only significantly enriched by gene set enrichment analysis (GSEA) of the phosphoproteome in COX17 KO cells (NES = 1.99, p = 0.002) but not CTR1 KO cells (Figure 6C, NES = 1.18, p = 0.26). Similarly, principal component 1 of modified AMPK pathway phosphopeptides in both mutants revealed that AMPK signaling is increased in COX17 compared to CTR1 KO cells (Figure 6I, p < 0.0001). This overall increase in phosphorylation of AMPK targets specifically in COX17 KO cells is consistent with their increased phosphorylation of PFKP (see Figure 3), which has been shown to directly interact with AMPK, an interaction that is enhanced by glucose starvation and promotes mitochondrial recruitment of AMPK (Chen *et al*., 2022). Notably, CTR1 KO cells had reduced phosphorylation of PFKP at S386 (0.56x wild-type levels, Supplemental Data File 1). This led us to hypothesize that preferentially increased AMPK activity inhibits mTOR in COX17 KO but not CTR1 KO cells, which we predicted should blunt mTOR-S6K activity relative to CTR1 KO cells (Figure 6S).

To explore mTOR-S6K activity, we looked at the abundance and phosphorylation of key mTOR-S6K pathway proteins in our proteomics dataset. The abundance of DEPTOR, an mTOR inhibitor (Peterson *et al*., 2009), and EIF2AK3 (PERK, eukaryotic translation initiation factor 2 alpha kinase 3), an ER stress response kinase that acts as a negative regulator of protein synthesis (Almeida *et al*., 2022), was reduced in both mutants but was significantly lower in CTR1 KO as compared to COX17 KO cells (Figure 7A). We confirmed these decreases in abundance by immunoblot (Figure 7D-E, (Lane *et al*., 2025b)). Levels of RPS6KA6, a member of the ribosomal S6K (RSK) family, were significantly lower in COX17-deficient cells as compared to CTR1-deficient cells (Figure 7A). While mTOR, AKT1S1, and RAPTOR had increased phosphorylation in both mutants as compared to wild-type cells, we observed greater phosphorylation of proteins downstream of mTOR signaling in CTR1-null cells compared to COX17-null cells, including EIF4G1 (eukaryotic translation initiation factor 4 gamma 1) at S984 (Raught *et al*., 2000) and multiple sites of RPS6 (Holz *et al*., 2005; Roux *et al*., 2007; Magnuson *et al*., 2012; Meyuhas, 2015) (Figure 7B). Principal component 1 of mTOR substrates in both mutants revealed that mTOR signaling is enriched in CTR1 KO but not COX17 KO cells (Figure 7C, p < 0.0001). We observed a small decrease in RAPTOR by immunoblot but observed no change in abundance by TMT-MS, and we detect no change in levels of RICTOR (Figure 7A, 7D-E).

**Figure 7.**
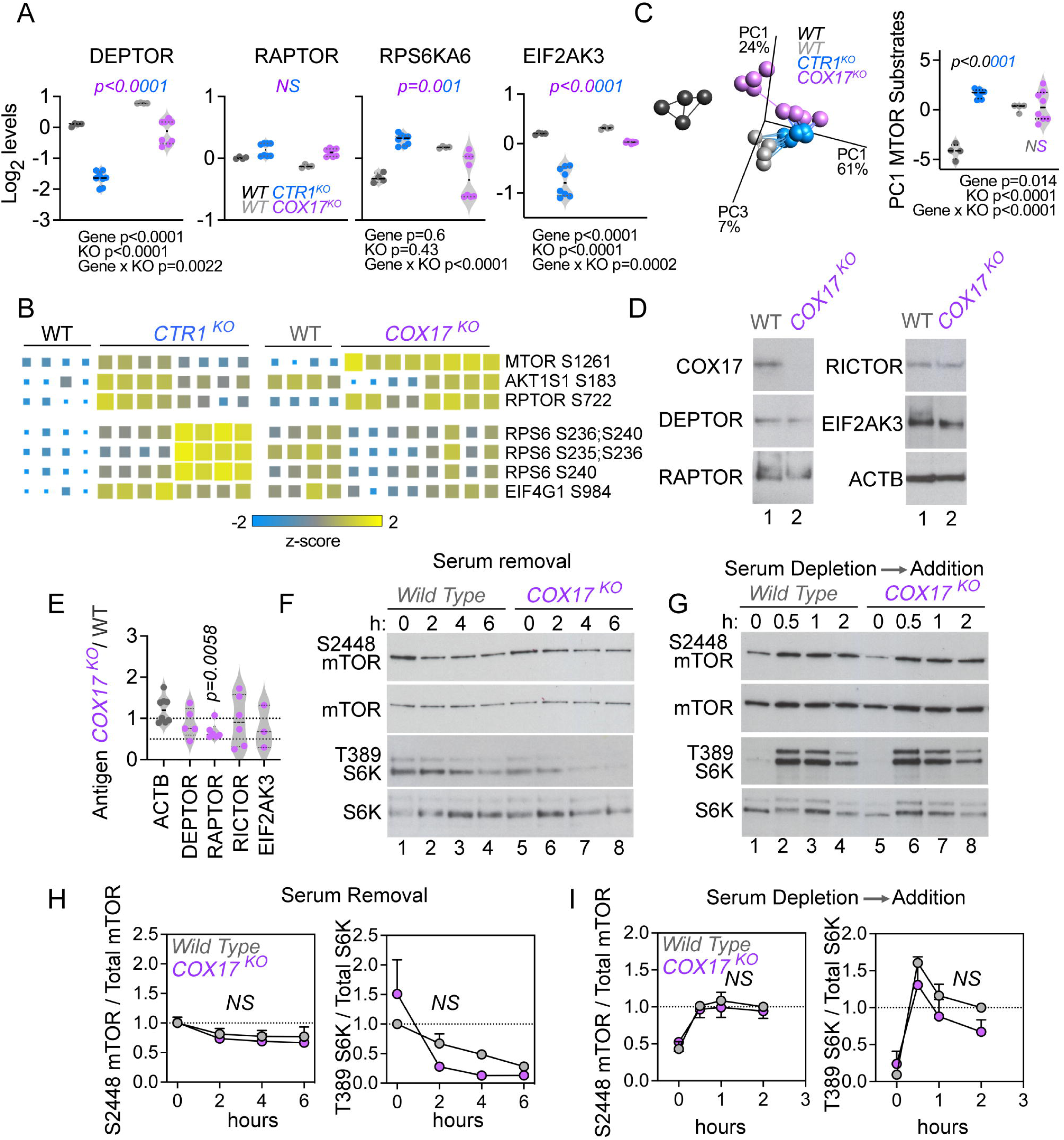
Reduced mTOR-S6K pathway activity in COX17-null cells as compared to CTR1-null cells. **A.** Mass spectrometry quantification of ontologically selected proteins (two-way ANOVA followed by Bonferroni’s multiple comparisons test) from two CTR1 KO clonal lines (blue symbols), two COX17 KO clonal lines (purple symbols), and their corresponding wild type clonal lines (dark gray and light gray, respectively). See Supplemental Data File 1 for source data. **B.** Heat map of mTOR pathway annotated phosphoproteins whose differential expression is significant with q < 0.05 (one-way ANOVA followed by Benjamini-Hochberg FDR correction). Each box represents a replicate. **C**. Principal component analysis (PCA) of mTOR substrates in **B** and PCA 1 (PC1) of the heat map entries comparing isogenic wild type, two COX17 KO clonal lines, and two CTR1 KO clonal lines. Two-way ANOVA followed by Bonferroni’s multiple comparisons tests. **D-E**. Immunoblots of whole cell extracts from wild type and COX17 mutant cells probed for COX17, DEPTOR, RAPTOR, RICTOR, EIF2AK3, and ACTB as a loading control. Immunoblots were quantified by normalizing protein abundance to wild-type cells. Italicized numbers represent p values analyzed by unpaired two-sided permutation t-test. **F-I**. Immunoblots of phosphorylated or total mTOR (pS2448) or S6K (pT389) from whole cell extracts from wild type and COX17 mutant cells at time 0 followed removal of fetal bovine serum for 2-6 hours (**F** and **H**) or after overnight depletion of fetal bovine serum followed by serum addition for 0.5-2 hours (**H** and **I**). Graphs depict quantitation of the ratio of the phosphorylated to total protein content, normalized to control at time 0 (**H**) or time at 2 hours (**I**) (n = 3-6 independent replicates per genotype, two-way ANOVA followed by Benjamini, Krieger, and Yekutiel corrections). CTR1 KO cell data in panels **A-C** was obtained concurrently and was reported in Lane *et al*. (2025b).

We further probed mTOR-S6K activity by manipulating serum levels in the media of COX17-deficient cells (Figure 7F-I), as activity of mTOR-S6K and PI3K-Akt signaling pathways can be stimulated by serum addition or inhibited by serum removal (Liu and Sabatini, 2020). As readouts, we measured the phosphorylation of mTOR at S2448 and S6K1 at T389, sites associated with increased activity of this pathway (Burnett *et al*., 1998; Navé *et al*., 1999; Chiang and Abraham, 2005). We previously used this paradigm to demonstrate heightened mTOR-S6K activity in CTR1 KO cells (Lane *et al*., 2025b). While serum removal decreased phosphorylation of both mTOR and S6K (Figure 7F,H) and overnight serum depletion followed by serum addition increased phosphorylation of both mTOR and S6K (Figure 7G,I), we observed no difference between the responses of COX17 KO and wild-type cells (Figure 7F-H; compare lanes 2-4 and 6-8). This is in marked contrast to CTR1 KO cells, which exhibited increased phosphorylation of both mTOR and S6K at baseline that was resistant to serum removal (Lane *et al*., 2025b).

Collectively, these data demonstrate that elevated mTOR-S6K activity is specific to CTR1 KO cells and is not observed in COX17-deficient cells (Figure 6S). We speculate that mTOR inhibition due to increased AMPK activation specifically in COX17 KO cells underlies the blunted mTOR-S6K activity in COX17-deficient cells as compared to CTR1 KO cells (note the highest ranked genes by GSEA, Figure 6C, and patterns of phosphorylation upstream and downstream of AMPK in the heat map, Figure 6H).

### Compartmentalized copper dyshomeostasis determines the requirement of mTOR for survival

To interrogate the role of mTOR signaling in the survival and growth of COX17 KO and CTR1 KO cells, we pharmacologically manipulated mTOR activity and quantitatively assessed the effects in cell survival assays (Figure 8), as we previously reported in CTR1 KO cells (Lane *et al*., 2025b). We first measured survival with increasing concentrations of serum in the media. We previously reported that CTR1 KO cell survival is resistant to serum removal (Lane *et al*., 2025b), and we predicted that we would observe the opposite phenotype in COX17 KO cells, given their reduced levels of mTOR signaling. As expected, COX17 KO cells survival under conditions of low serum was reduced as compared to both wild type and CTR1 KO cells (Figure 8A), suggesting that COX17 deficiency makes cell survival more susceptible to mTOR inhibition.

**Figure 8.**
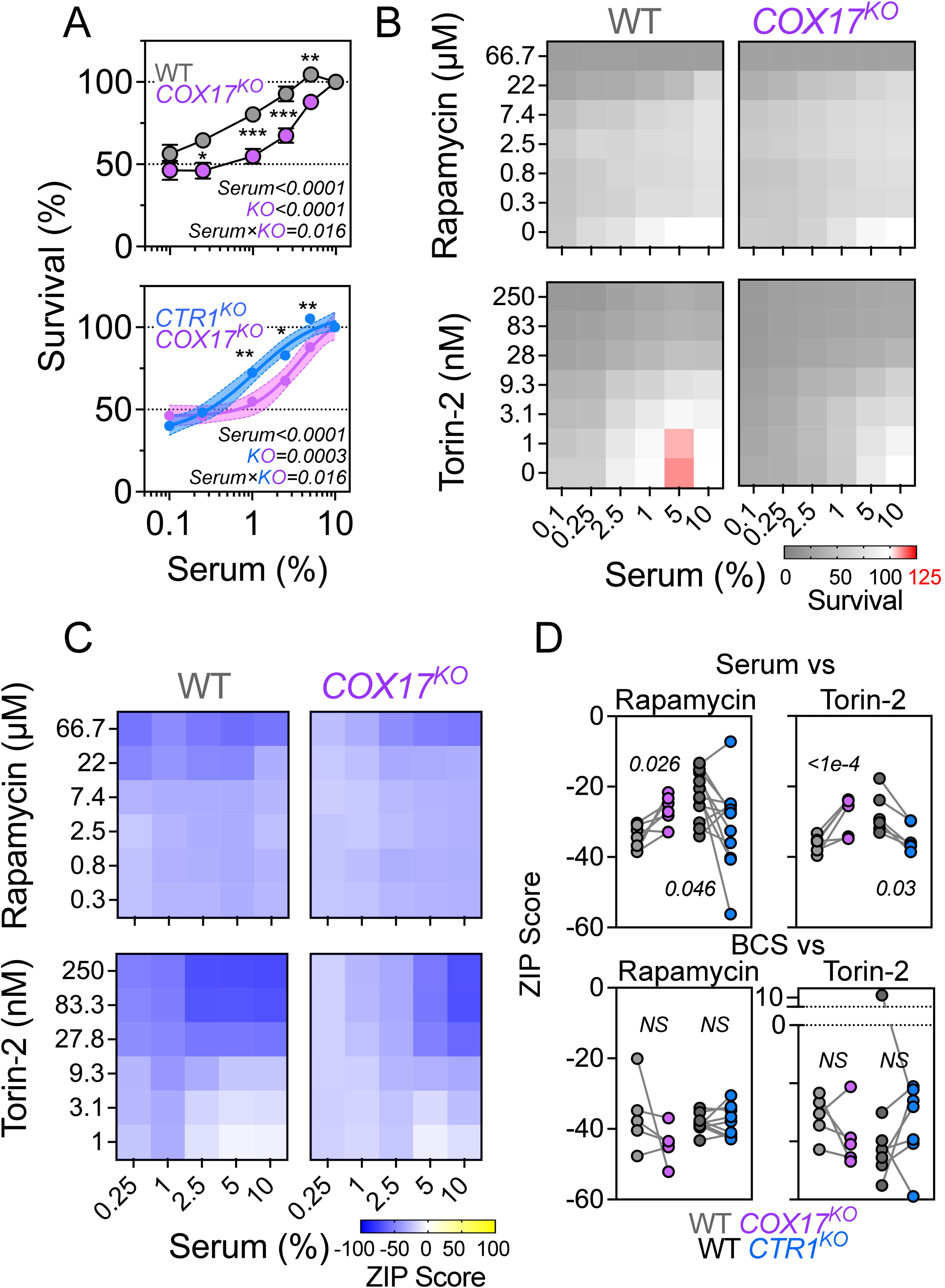
Inactivation of the mTOR pathway is adaptive in COX17-null cells. **A.** Cell survival analysis of wild type and COX17 mutants with increasing concentrations of serum (top; average ± SEM, n *=* 13) or comparing isogenic COX17 and CTR1 KO mutants (bottom; average ± 95% CI, n *=* 13 for COX17 or n = 18 for CTR1 mutants). Two-way ANOVA followed by Bonferroni’s multiple comparisons test. **B-D.** Synergy analysis of cell survival of isogenic COX17 and CTR1 mutants treated with combinations of serum, rapamycin, Torin-2, and BCS at increasing concentrations. **B.** Cell survival map for cells treated with serum and the mTOR inhibitors rapamycin or Torin2 (n = ≥ 5 independent experiments per pair), with the corresponding interaction synergy map (**C**) calculated using the Zero Interaction Potency (ZIP) score for cell survival (Yadav *et al*., 2015). Scores below −10 indicate an antagonistic interaction between the compounds. **D**. Average ZIP score for drug interactions between either rapamycin or Torin2 and serum or BCS (average ± SEM, paired estimation statistics). CTR1 KO cell data in panels **A** and **D** was obtained concurrently and was reported in Lane *et al*. (2025b).

Next, we performed cell survival assays with different combinations of mTOR-specific inhibitors and the copper chelator BCS to determine how these compounds interacted synergistically or antagonistically to affect cell survival using the zero-interaction potency (ZIP) model. The ZIP model assumes no interaction between drugs (a ZIP score of 0) and predicts an interaction is likely to be antagonistic if the ZIP score is less than −10 (Yadav *et al*., 2015; Ianevski *et al*., 2022). We first tested the effect of the combination of serum and the mTOR inhibitors rapamycin and Torin-2 on COX17 KO survival. Torin-2 inhibits both mTORC1 and mTORC2, while rapamycin is the canonical mTORC1 inhibitor (Ballou and Lin, 2008; Zheng and Jiang, 2015). We observed antagonistic responses between serum and both rapamycin and Torin-2 across all genotypes (Figure 8B-D), indicating that the mTOR inhibitors counteract pro-survival effects of serum. However, the effect of mTOR inhibition differed depending on the mutated gene. mTOR inhibition made the interaction between serum and rapamycin or Torin-2 in COX17-deficient cells *less* antagonistic, indicating a heightened survival of COX17 cells in the presence of mTOR inhibitors, while the opposite was observed in CTR1-deficient cells (Figure 8D). There were no genotype-dependent effects on survival by the combination of the copper chelator BCS and rapamycin or Torin-2 (Figure 8D). We interpret these results to indicate that mTOR activity is a pro-survival adaptive response in CTR1-deficient but not COX17-deficient cells.

To explore the role of mTOR pathway activity with respect to compartmentalized copper deficiency in a whole organism, we studied the effect of S6K RNAi (S6K-IR) on the survival of models of cellular copper deficiency (ATP7 overexpression, ATP7-OE) and COX17 deficiency (COX17-IR) in the *Drosophila* epidermis (Fig. 9, Fig. S9). ATP7-OE is a highly validated model of cell-autonomous copper deficiency due to enhanced metal efflux (Norgate *et al*., 2006; Binks *et al*., 2010; Hwang *et al*., 2014; Hartwig *et al*., 2020; Lane *et al*., 2025b). We drove the expression of UAS-RNAi or UAS-ATP7 transgenes with the trans activator pnr-GAL4, which restricts expression to epidermal epithelium (Norgate *et al*., 2006; Binks *et al*., 2010; Hwang *et al*., 2014; Hartwig *et al*., 2020; Lane *et al*., 2025b). We assessed the impact of these genotypes across sexes by scoring the hazard ratio, a metric that describes the risk of an event occurring over time compared to another group (Breslow, 1975). A hazard ratio >1 indicates increased risk of death while a ratio <1 indicates decreased death risk. Copper deficiency arising from ATP7-OE alone decreased survival in both sexes (Fig. 9A,C and Fig. S9A,C), with a hazard ratio of 1.15 in males and 2.28 in females (Fig. 9B and Fig. S9B). In contrast, COX17 deficiency and S6K deficiency had opposite effects on survival depending on sex. COX17-IR led to increased survival in males (Fig 9A, hazard ratio of 0.67, FDR 1.12e-10) and decreased survival in females as compared to with type animals (Fig S9, hazard ratio of 1.37, FDR 0). S6K-IR increased survival in males and decreased survival in females (Fig. 9 and S9, 0.75 and 1.14, FDR 0.0014 and 0.015). This sexual dimorphic phenotype has been previously described in fly mitochondrial mutants, with female *Drosophila* exhibiting increased resilience to genetic defects affecting mitochondria (Binks *et al*., 2010; Innocenti *et al*., 2011; Hwang *et al*., 2014; Kemppainen *et al*., 2014; Patel *et al*., 2016). Therefore, we focused on male survival in analysis of combinatorial genetic defects, as this sex displayed the ATP7-OE and COX17-IR hazard ratios closest to wild type. We reasoned this could avoid potential flooring effect on survival when combined with other transgenes. Reducing the expression of S6K decreased the survival of ATP7-OE males (Fig. 9, compare ATP7-OE and ATP7-OE/S6K-IR, hazard ratios of 1.17 and 1.7, FDR 0.0022). In contrast, decreased levels of S6K did not significantly affect the survival of COX17 deficient male flies (Fig. 9, compare COX17-IR and COX17-IR/S6K-IR, hazard ratios of 0.67 and 0.76, FDR 0.58). The effects of these transgene combinations were attenuated or absent in female flies (Fig. 9S). This suggests that S6K expression is dispensable for survival in COX17-deficient but not copper deficient ATP7-OE animals.

**Figure 9.**
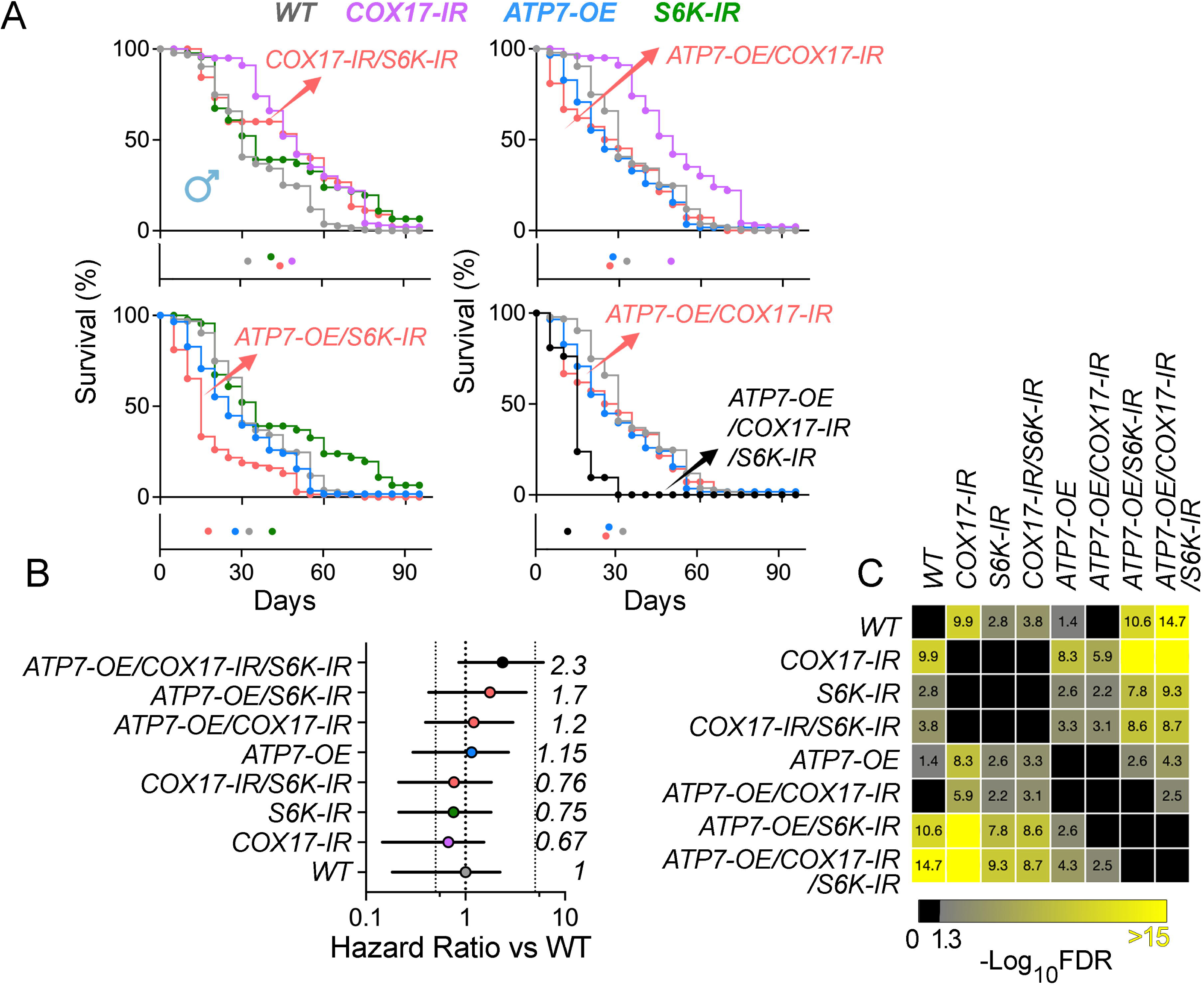
Male survival and hazard structure in copper dyshomeostasis-S6K interaction genotypes. **A.** Kaplan–Meier survival curves for males. Eight genotypes were tested: *pnr_W1118* (WT reference, n=187), *pnr_COX17_RNAi_111465* (COX17-IR, n=100), *pnr_S6K_RNAi* (S6K-IR, n=46), the combined *pnr_COX17_111465_S6K_RNAi* double knockdown (COX17-IR/SK6-IR, n=45), ATP7 overexpression (ATP7wt_pnr_W1118, ATP7-OE, n=58), and ATP7 overexpression combined with COX17 (n=28) and/or S6K (n=69 and 21 or triple transgenic) knockdown. Survival was reconstructed from percent-alive data and starting cohort size per genotype. Male Kaplan–Meier curves show strong survival loss following COX17 and S6K reduction, with additive or greater-than-additive decline in double and ATP7-sensitized backgrounds. **B.** Hazard ratio (HR) forest plots relative to WT. Per-genotype hazards were estimated using death rate per animal-day, normalized to WT hazard. Points represent HR, error bars indicate 95% CIs. **C.** Pairwise log-rank comparison heatmaps (Benjamini–Hochberg FDR). Matrix shows - log_10_(FDR) for all pairwise genotype survival.

Next, we asked whether phenotypes caused by reduced levels of S6K could become overt in both sexes where cellular and mitochondrial copper content were simultaneously decreased. We hypothesized that the expressivity of the mortality phenotype observed in either ATP7-OE or ATP7-OE/S6K-IR double transgenic flies should be increased in animals mimicking a more severe copper deficiency simultaneously impairing copper homeostasis in whole cells (ATP7-OE) and mitochondria (COX17-IR). Triple transgenic flies (ATP7-OE/COX17-IR/S6K-IR) displayed the most severe mortality phenotype, with an increased risk of death 2.3-fold higher than wild type males and 1.91 times higher than ATP7-OE/COX17-IR in males (Fig. 9, FDR 2.07e-15 and 0.0031). The requirement of S6K in cells with altered copper homeostasis also became evident in triple transgenic female flies (ATP7-OE/COX17-IR/S6K-IR) as compared to ATP7-OE/COX17-IR. The risk of death was increased by 1.76-fold as compared to ATP7-OE/COX17-IR females (Fig. S9, FDR 0.015).

These results show that S6K is necessary for survival in cellular copper deficiency (ATP7-OE) but is dispensable for survival in a model of mitochondrial copper dyshomeostasis (COX17-IR) in a sex-dependent manner. We conclude that S6K is dispensable or deleterious in compartmentalized mitochondrial models of copper deficiency while protective in models that deplete cells of copper.

## Discussion

Here, we report both unique and shared mechanisms downstream of compartmentalized copper deficiency by selectively targeting copper delivery to complex IV in the mitochondrion in COX17 mutants or by disrupting cellular copper homeostasis globally in CTR1 mutants. We concurrently generated and performed proteomics and transcriptomic studies in these isogenic COX17 KO, CTR1 KO, and wild-type clones, thus allowing direct comparisons of molecular phenotypes between these genotypes (Figure 1, Lane *et al*. (2025b)). CTR1- and COX17-null cells share select phenotypes, including mitochondrial copper deficiency, increased glycolysis, and reduced levels and impaired assembly of complex IV and other respiratory complexes, with more dramatic respiratory defects in COX17-null cells (Figure 2), but exhibit marked differences in the identity of genes and proteins affected by the mutation. Many proteins that were commonly affected by both genotypes differed in the magnitude and/or direction of change in abundance, including proteins belonging to the mitochondrion, the actin cytoskeleton, the Golgi complex, the lysosome, and the nucleus as well as the mTOR and AMPK signaling pathways, representing unique mechanisms engaged by each mutant despite their shared defect in mitochondrial copper content.

Distinct molecular mechanisms observed in COX17 but not CTR1 KO cells include reduced abundance and phosphorylation of multiple chromosomal passenger complex (CPC) and THO complex components (Figure 3, Supplemental Data File 1), as well as upregulation of genes involved in DNA repair and response to DNA damage, including ERCC6, BRCA2, ARIDS2, MSH2, and NRDE2 (Supplemental Data File 1). Additionally, multiple cytoskeletal proteins exhibit opposite patterns of abundance and/or phosphorylation in each mutant, with ABLIM1, AHNAK, MYLK, and TAGLN increasing in abundance and/or phosphorylation in COX17 KO cells and decreasing in CTR1 KO cells; multiple proteins involved in cadherin and actin binding follow a similar pattern (Figure 3S1, Supplemental Data File 1). PERK (*EIF2AK3*), a copper-dependent kinase (Bond Newton *et al*., 2025), exhibits decreased abundance and phosphorylation (S555) in CTR1 KO but not COX17 KO cells (Supplemental Data File 1) and may be a key regulator of these cytoskeleton and cell adhesion phenotypes (Diaz *et al*., 2025). With respect to the mitochondria, the specific enrichment in mitochondrial membrane proteins in COX17-deficient cells but not CTR1-deficient cells is consistent with the proposed role of COX17 in the mitochondrial contact site and cristae organizing system (MICOS) and predicts alterations in the mitochondrial inner membrane fusion-fission cycle in COX17-null cells. COX17 directly interacts with the MICOS subunit Mic60, and this interaction is required for MICOS complex integrity (Chojnacka *et al*., 2015; Kozjak-Pavlovic, 2017). These specific examples highlight the broad differences in nuclear and cytoskeletal phenotypes in COX17 and CTR1 deficiency and predict disruption of the mitochondrial inner membrane and fusion-fission machinery in COX17-null cells.

Lysosomal changes were apparent in both mutants but were more exaggerated in COX17-null cells. While HEXA/B were upregulated in both mutants, this effect was more dramatic and only corresponded to increased abundance at the protein level in COX17-deficient cells (Figure 3, Figure 5, Supplemental Data File 1). This same pattern was observed for multiple cathepsins; increased levels of mature cathepsin B and D in COX17 KO cells were confirmed by immunoblot (Figure 4). We speculate that changes in the levels of lysosomal hydrolases may be linked to one or more of the following mechanisms: modifications in the sorting of lysosomal hydrolases from the Golgi complex to the lysosome, a compensatory process for autophagy defects, and/or AMPK-dependent mitochondria-lysosome signaling (Fernández-Mosquera *et al*., 2017; Fernandez-Mosquera *et al*., 2019; Deus *et al*., 2020). We predict impaired autophagy in both COX17 and CTR1 mutant cells. Impaired autophagy can be induced by promoting a hyperglycolytic state in neurons (Jimenez-Blasco *et al*., 2024), consistent with the increased glycolysis we observed in both mutants (Figure 2), or by mTOR-AMPK activity (Kim and Guan, 2015; Park *et al*., 2023), which we demonstrate is differentially impacted by COX17 and CTR1 mutations. Additionally, distinct autophagy mechanisms related to copper levels are likely engaged in CTR1 KO cells, as copper is required for kinase activity of ULK1/2 (Tsang *et al*., 2020), important regulators of autophagy initiation (Chan *et al*., 2007; Zachari and Ganley, 2017). Thus, we propose that a constellation of factors may ultimately result in increased content of lysosomal hydrolases and impaired autophagy through partially overlapping and partially distinct mechanisms in COX17 and CTR1 deficiency.

We identified the AMPK and mTOR pathways as key regulators of metabolism differentially impacted by COX17 and CTR1 KO (Figure 6S). A central difference between the two mutants is the pronounced activation in the AMPK pathway in COX17-null cells at the level of the transcriptome, proteome, and phosphoproteome with comparatively reduced activity of mTOR relative to CTR1-null cells, supported by multipronged evidence (Figures 5-7, 3S1, 5S). At the transcript level, changes in expression of AMPK pathway gene levels were significantly more pronounced in COX17 KO cells than CTR1 KO cells, including upregulation of three AMPK subunits (Figure 5). Increased total AMPKα1/PRKAA1 protein was observed in both mutants but was more pronounced in COX17 KO cells (1.54x wild-type levels in COX17 KO, 1.17x wild-type levels in CTR1 KO by immunoblot), and the same pattern emerged for AMPK phosphorylation and direct substrates of AMPK (Figure 6, Supplemental Data File 1) (Mihaylova and Shaw, 2011; Schaffer *et al*., 2015). This increased abundance and phosphorylation of AMPK in COX17 KO cells produced elevated phosphorylation of AMPK substrates not observed in CTR1 cells, such as eEF2, GAPVD1, and FOXO3 (Figure 3, Figure 6, Supplemental Data File 1; (Greer *et al*., 2007; Schaffer *et al*., 2015; Johanns *et al*., 2017; Wang *et al*., 2017b; Ducommun *et al*., 2019)). Increased eEF2 phosphorylation in COX17 cells is consistent with increased AMPK activity and reduced mTOR activity compared to CTR1 KO cells (Browne *et al*., 2004; Wang *et al*., 2014; Johanns *et al*., 2017; Kumar *et al*., 2020), which may be protective against nutrient deprivation (Leprivier *et al*., 2013; Yamada *et al*., 2019). Evidence in support of reduced mTOR activity in COX17 cells comes from the enrichment and more substantial alteration of mTOR-S6K pathway proteins in the proteome of CTR1 KO cells as compared to COX17 cells, such as increased phosphorylation of EIF4G1 (Raught *et al*., 2000) and RPS6 (Holz *et al*., 2005; Roux *et al*., 2007; Magnuson *et al*., 2012; Meyuhas, 2015) (Figure 3, Figure 6, Figure 6S Supplemental Data File 1). Accordingly, increased mTOR activity was identified in CTR1 KO but not COX17 KO cells by Kinase-Substrate Enrichment Analysis (KSEA), and principal component 1 analysis revealed that mTOR signaling is enriched specifically in CTR1 KO cells and not in COX17 KO cells (Figure 6, Figure 7). Additionally, COX17 cells did not exhibit elevated mTOR-S6K phosphorylation at baseline or under conditions of serum addition or removal that we previously reported in CTR1 KO cells (Lane *et al*., 2025b) (Figure 7). Further supporting reduced mTOR activity in COX17-deficient cells is their reduced phosphorylation of LARP1 (Supplemental Data File 1), which stabilizes the mTOR transcript (Mura *et al*., 2015; Jia *et al*., 2021), and decreased phosphorylation of the mTOR-S6K interactor EIF3F (Supplemental Data File 1, (Holz *et al*., 2005; Harris *et al*., 2006; An *et al*., 2020)). Central metabolic checkpoints that determine the activity of the mTOR cascade by AMPK are the phosphorylation of RAPTOR in serines 722 and 792 and TSC2 in residues 1227 and 1345 (Inoki *et al*., 2003; Gwinn *et al*., 2008). We did not find differential phosphorylation of these sites between COX17 and CTR1 KO cells. Thus, it is likely that other AMPK-dependent phosphorylation events along the mTOR pathway determine the protective effect mTOR inhibition of COX17 null cells as compared to CTR1 KO cells. As AMPK is the most upstream signaling hub in the KEGG mTOR and protein synthesis pathways (Figure 6S), these data support our contention that increased AMPK signaling in COX17 KO cells as compared to CTR1 KO cells contributes to blunted mTOR pathway activation observed solely in COX17 deficiency (see Figure 6S).

Another mechanism by which mTOR activity is blunted in COX17 KO cells by AMPK is FOXO3 signaling. COX17-null cells exhibit increased abundance and AMPK-and Akt-mediated phosphorylation of FOXO3 (Greer *et al*., 2007; Li *et al*., 2009) (see Figure 6S), which can block mTOR activity (Chen *et al*., 2010) or causes FOXO3 retention in the cytoplasm (Brunet *et al*., 1999; Plas and Thompson, 2003; Dobson *et al*., 2011). The FoxO family are important regulators of metabolism (Rodriguez-Colman *et al*., 2024), and FOXO3 regulates mitochondrial membrane potential, respiration, mitochondrial architecture, energy metabolism, and cellular ROS in neuronal tumor cells (Hagenbuchner and Ausserlechner, 2013). Thus, FOXO3 may represent a key node in a mitochondrial signaling hub in COX17 deficiency. Future studies can further explore the mechanism responsible for FOXO3 activity, but we speculate that COX17 cells exhibit increased p53 activity that could underly this FOXO3 phenotype and other phenotypes reported here. FOXO3 is a p53 target gene (Renault *et al*., 2011) and plays a role in responding to DNA damage (Chung *et al*., 2012). TAGLN, which is elevated in COX17 KO cells, can also be induced by p53 (Tsui *et al*., 2019). An additional 18 proteins identified in a meta-analysis as p53 target genes (Fischer, 2017) increase in abundance in COX17 cells relative to wildtype, including MYLK, STEAP3, and TP53I3. p53 pathway activation inhibits mTOR and activates AMPK (Budanov and Karin, 2008; Kon *et al*., 2021). Thus, we propose that the convergence of increased AMPK, FOXO3, and p53 pathway activation blunts mTOR activation in response to COX17 but not CTR1 deficiency, such that CTR1 KO cells retain increased mTOR-S6K activity.

The functional impact of this difference in mTOR and S6K activity between COX17 and CTR1 KO cells was demonstrated by the differential effects of mTOR activity on cell survival. In CTR1-deficient cells, mTOR *activation* is prosurvival whereas in COX17 KO cells, mTOR *inhibition* is prosurvival (Figure 8). We previously reported that increased mTORC1/S6K activation is an adaptive and prosurvival mechanism in CTR1 KO cells and *Drosophila* models of neuronal copper deficiency (Lane *et al*., 2025b). These results are also supported by a recent preprint in which prenatal pharmacological mTOR activation in a copper chelation model of copper deficiency rescued oligodendrocyte developmental defects and brain weight during embryogenesis and improved social deficits in the adult animal (Usui *et al*., 2023). Our findings in mammalian cells are conserved in *Drosophila*, where S6K is necessary for survival in animals with cell autonomous copper deficiency (ATP7-OE males), akin to CTR1 KO in cell lines, as well as more severe copper dyshomeostasis animals with simultaneous disruption of both cellular and mitochondrial copper homeostasis (ATP7-OE/COX17-IR, both males and females). In stark contrast, the observation that mTOR inhibition is adaptive in COX17-deficient cells corroborates the established prosurvival effects of mTOR inhibition in mouse models of mitochondrial diseases (Johnson *et al*., 2013; Siegmund *et al*., 2017). This effect is also observed in vivo, as S6K is dispensable for survival in a model of mitochondrial copper dyshomeostasis induced by COX17 RNAi. We conclude that mTOR signaling and downstream S6K activity plays a conserved role in copper homeostasis: S6K activity is either dispensable or deleterious in compartmentalized mitochondrial models of copper deficiency but is protective in cellular copper deficiency (Lane *et al*., 2025b).

Finally, our studies also revealed three intriguing observations that reflect the complexity of the mechanisms controlled by compartment-specific copper homeostasis and inform our understanding of COX17 function, which has received comparatively little study as compared to other key cuproproteins (Lutsenko *et al*., 2024; Lane *et al*., 2025a). First, it is noteworthy that mitochondrial copper levels were reduced to a similar degree in both CTR1 and COX17 mutants. The cellular machinery responsible for loading copper into the mitochondria and for assembly of complex IV has undergone extensive study but remains only partially understood (Cobine *et al*., 2006; Bourens *et al*., 2013; Boulet *et al*., 2018; Cobine *et al*., 2021; McCann *et al*., 2022; Lutsenko *et al*., 2024). COX17 is present in both the cytosol and mitochondrial intermembrane space (Beers *et al*., 1997; Heaton *et al*., 2000), and it remains unknown whether COX17 acquires copper within the mitochondrial matrix or delivers copper into the mitochondria from the cytosol (Cobine *et al*., 2004). Inner mitochondrial membrane COX17 expression is sufficient to support complex IV activity (Maxfield *et al*., 2004), suggesting that COX17 can acquire copper within the matrix for delivery to complex IV; however, the reduction in mitochondrial copper we observed in COX17-null cells, consistent with a recent report characterizing mutant SCO1 cells with impaired COX assembly (Ghosh *et al*., 2025), suggests that COX17 and other complex IV assembly factors and chaperones play an important role in overall mitochondrial copper homeostasis. Second, metal content differed between COX17 and CTR1 mutants. Copper was the only metal with altered levels in CTR1 KO cells, while COX17 KO mitochondria had increased iron and decreased potassium (1.72x and 0.63x wildtype levels, respectively; Figure 1). While speculative, the increase in iron in the mitochondria of COX17 KO cells could originate in respiration-dependent mitochondrial redox imbalances more pronounced in COX17 KO cells than CTR1 KO cells (Chen *et al*., 2023). Interestingly, a recent report indicated that ablation of the copper transporter *atp7a* induced iron overload in zebrafish larvae, and axonal and myelin defects in *atp7a*-null animals were rescued by iron chelation of inhibiting ferroptosis (Wu *et al*., 2025). Together, this suggests the intriguing idea that iron phenotypes in *atp7a*-deficient animals could be downstream of pronounced mitochondrial copper deficiency and reduced complex IV activity observed in advanced stages of Menkes disease, which may be emulated by COX17 deficiency.

Third, we observed an unexpected drastic reduction in abundance of the secretory copper-dependent enzyme DBH in COX17-null as compared to CTR1-null cells (Figure 4). We argue that this reduction in DBH levels in COX17 vs. CTR1 KO cells is not dependent on total and mitochondrial copper levels, as total copper is unchanged in COX17 KO cells and both mutants have similar levels of mitochondrial copper. Neither is this DBH phenotype representative of a global secretory defect in COX17 KO cells, as their levels of the secretory protein chromogranin B are normal (Figure 4) and we previously reported that COX17-deficient cells increase APOE content and secretion by 15-fold (Wynne *et al*., 2023). We propose that COX17 is required for DBH biosynthesis, either directly or through mechanisms involving ATP7A. COX17 lacks a classic mitochondrial import sequence (Diekert *et al*., 1999) and is targeted to the mitochondria by a Mia40-Erv1 disulfide relay system in the mitochondrial intermembrane space (Mesecke *et al*., 2005; Bragoszewski *et al*., 2013; Chojnacka *et al*., 2015). Thus, COX17 possesses unique characteristics in terms of its intracellular trafficking. In fact, 56% of the COX17 pool is localized to the cytoplasm, in contrast to other intermembrane proteins whose cytoplasmic levels are under 7% (Itzhak *et al*., 2016), yet additional cytosolic roles for COX17 remain to be explored. COX17 has been proposed to be a natively unfolded protein that acts as a shuttle between the cytoplasm and the mitochondria (Punter and Glerum, 2003). This proposed function is based on its dual localization in the cytoplasm and mitochondria and similarities in its amino acid sequence to the natively unfolded protein Nup2, which traffics between the nucleoplasm and cytoplasm (Denning *et al*., 2002). The intriguing possibility that such a shuttling capacity is required for appropriate copper entry to the Golgi through ATP7A-independent or -dependent mechanisms remains to be studied.

Together, these data suggest that both CTR1- and COX17-null cells could serve as cellular models with which to study compartmentalized copper deficiency and Menkes disease mechanisms. Our observations of distinct molecular mechanisms engaged in both mutants, particularly altered AMPK-mTOR signaling, may reflect the progression of Menkes disease from a “CTR1 KO-like” to a “COX17 KO-like” phenotype as the brain continues to suffer severely reduced or absent complex IV activity. Early mTOR-S6K activation and stimulation of protein synthesis may be temporarily protective, consistent with the prosurvival effects of mTOR activation in CTR1 KO cells and the rescue of dendritic phenotypes in *Drosophila* models of copper deficiency following genetic stimulation of mTOR-S6K pathway activity (Lane *et al*., 2025b); however, mTOR activity may ultimately become maladaptive over time as Menkes disease and neurodevelopment progresses, reflecting the prosurvival effects of mTOR inhibition in COX17-deficient cells and animal models of mitochondrial diseases (Johnson *et al*., 2013; Siegmund *et al*., 2017).

## Materials and Methods

### Cell lines, gene editing, and culture conditions

Human neuroblastoma SH-SY5Y cells (ATCC, CRL-2266; RRID:CVCL_0019) were cultured and mutant lines generated as described in Lane *et al*. (2025b). Cells were grown in DMEM media (Corning, 10-013) containing 10% FBS (VWR, 97068-085) at 37°C in 5-10% CO_2_. COX17-deficient cells were generated by Synthego by genome editing using gRNA and Cas9 preassembled complexes with a knock-out efficiency of 85% with the gRNAs CCAAGAAGGCGCGCGAUGCG, which targeted transcript ENST00000261070 at exon 1. Clonal cell lines were generated from wild-type and mutant pools by limited dilution. Sanger sequencing was used to confirm mutagenesis with the primer: 5’ AGGCCCAATAATTATCTCCAGAGC. All controls represent either a single wild-type clone or a combination of two wild-type clones. All experiments used two separate mutant clones of cells (KO3 and KO5) to exclude clonal or off-target effects unless otherwise indicated. CTR1-deficient SH-SY5Y cells were generated and clones isolated as described in Lane *et al*. (2025b).

### Antibodies

Table 1 lists the antibodies used at the indicated concentrations for western blots and immunofluorescence.

**Table 1.** Antibodies.

| Antibody | Dilution | Catalog Number | RRID |
| --- | --- | --- | --- |
| Actin B | 1:5000 | Sigma-Aldrich A5441 | AB_476744 |
| AMPK | 1:1000 | Cell Signaling 2532 | AB_330331 |
| AMPK P-T172 | 1:1000 | Cell Signaling 2535 | AB_331250 |
| ATP7A | 1:500 | NeuroMab 75-142 | AB_10672736 |
| Cathepsin B | 1:1000 | ProteinTech 12216-1-AP | AB_2086929 |
| Cathepsin D | 1:5000 | ProteinTech 21327-1-AP | AB_10733646 |
| CCS | 1:500 | ProteinTech 22802-1-AP | AB_2879172 |
| Chromogranin B (CgB) | 1:500 | ProteinTech 14968-1-AP | AB_2081138 |
| COX17 | 1:500 | ProteinTech 11464-1-AP | AB_2085109 |
| COX4 | 1:1000 | Cell Signaling 4850 | AB_2085424 |
| CTR1 (SLC31A1) | 1:2000 | ProteinTech 67221-1-IG | AB_2919440 |
| DBH | 1:500 | Millipore AB1536 | AB_2089474 |
| DEPTOR | 1:1000 | Cell Signaling 11816 | AB_2750575 |
| FOXO3 | 1:1000 | Cell Signaling 12829 | AB_2636990 |
| GM130 | 1:250 | BD Biosciences 610822 | AB_398141 |
| HSP90 | 1:1000 | BD Biosciences 610418 | AB_397798 |
| Integrin α1/CD49a | 1:1000 | Cell Signaling 15574T | AB_2536530 |
| mTOR | 1:1000 | Cell Signaling 2983 | AB_2105622 |
| mTOR pSer2448 | 1:1000 | Cell Signaling 5536 | AB_10691552 |
| MYLK | 1:1000 | ProteinTech 21642-1-AP | AB_10734590 |
| NDUFB11 | 1:500 | Abcam ab183716 | AB_2927481 |
| OXPHOS mix | 1:250 | Abcam ab110412 | AB_2847807 |
| PERK (EIF2AK3) | 1:1000 | Cell Signaling 5683 | AB_10841299 |
| Raptor | 1:1000 | Cell Signaling 2280 | AB_2737590 |
| Rictor | 1:1000 | Cell Signaling 2114 | AB_561245 |
| S6K P70 | 1:1000 | Cell Signaling 9202 | AB_2179963 |
| S6K pThr389 | 1:1000 | Cell Signaling 9234 | AB_331676 |
| SDHA | 1:1000 | 11998 | AB_2269803 |
| Steap3 | 1:1000 | ProteinTech 28478-1-AP | AB_3086055 |
| TAGLN | 1:1000 | Cell Signaling 36090 | AB_3697810 |
| THOC2 | 1:2000 | Bethyl A303-630A | AB_11205629 |
| THOC5 | 1:2000 | Bethyl A302-119A | AB_1604294 |
| UQCRC2 | 1:500 | Abcam ab14745 | AB_2213640 |
| mouse HRP | 1:5000 | A10668 | AB_2213640 |
| rabbit HRP | 1:5000 | G21234 | AB_2534058 |

### Drugs

Table 2 lists the drugs used at the indicated concentrations or concentration ranges as described in their corresponding figure legends.

**Table 2.** Drugs.

| Drug | Source/Cat. No | Storage/Stock | Concentration range |
| --- | --- | --- | --- |
| BCS | Sigma, B1125 | 400 mM, DMSO, -20C | 0.1 mM - 1.6 mM |
| Oligomycin | Sigma, 75351 | 10 mM, DMSO, -20C | 1.0 μM |
| FCCP | Sigma, C2920 | 10 mM, DMSO, -20C | 0.25 μM |
| Rotenone | Sigma, R8875 | 10 mM, DMSO, -20C | 0.5 μM |
| Antimycin A | Sigma, A8674 | 10 mM, DMSO, -20C | 0.5 μM |
| D-Glucose | Sigma, G8769 | 2.5M, 4C | 10 mM |
| 2-deoxyglucose | Sigma, D3179-1G | 500 mM, Glycolysis Stress Test Media, -20C | 50 mM |
| Torin-2 | VWR, 103542-338 | 1 mM, DMSO, -20C | 1 nM - 250 nM |
| Rapamycin | VWR, 101762-276 | 50 mM, DMSO, -20C | 0.8 μM - 66 μM |

### Immunoblotting

Cells were collected and protein isolated as described in Lane *et al*. (2025b), and treatments are described in each figure. Briefly, cells were grown in 12 or 24 well plates. After reaching desired confluency, the plates were placed on ice and washed with cold phosphate-buffered saline (PBS) (Corning, 21-040-CV). Lysis buffer (Buffer A with 0.5% Triton X-100; Sigma, T9284) with Complete anti-protease (Roche, 11245200) was added to each plate. PhosSTOP phosphatase inhibitor (Roche, 04906837001) was also added to the lysis buffer for samples of interest for phosphorylated proteins. Buffer A is contains 150 mM NaCl, 10 mM HEPES, 1 mM ethylene glycol-bis(β-aminoethylether)-*N*,*N*,*N*′,*N*′-tetraacetic acid (EGTA), and 0.1 mM MgCl2, pH 7.4. Cells were scraped and placed in Eppendorf tubes, left on ice for 20 min, and centrifuged at 16,100 × *g* for 10 min and the clarified supernatant was recovered. The Bradford Assay (Bio-Rad, 5000006) was used to determine protein concentration, and all lysates were frozen and stored at −80C until use.

Immunoblotting was conducted as described previously (Lane *et al*., 2025b). To reduce and denature cell lysates, they were combined with Laemmli buffer (SDS and 2-mercaptoethanol) and heated for 5 min at 75C. Equivalent amounts of samples were loaded onto 4-20% Criterion gels (Bio-Rad, 5671094) for SDS-PAGE and transferred using the semidry transfer method to polyvinylidene difluoride (PVDF) membranes (Millipore, IPFL00010). The membranes were blocked in TRIS-Buffered Saline (TBS) containing 5% nonfat milk and 0.05% Triton X-100 (TBST) for 30-45 min at room temperature (RT). After rinsing the membranes thoroughly, the membranes were incubated with optimally diluted primary antibody in a buffer containing PBS with 3% bovine serum albumin (BSA) and 0.2% sodium azide overnight. Following rinses with TBST, membranes were incubated with horseradish peroxidase-conjugated secondary antibodies against mouse or rabbit (see Table 1) diluted 1:5000 in TBST with 5% nonfat milk for 30-60 min at RT and rinsed with TBST at least three times. Membranes were probed with Western Lightning Plus ECL reagent (PerkinElmer, NEL105001EA) and exposed to GE Healthcare Hyperfilm ECL (28906839). Additional staining was repeated as described above following stripping of blots (20 min with 200 mM glycine, 13.8 mM SDS, pH 2.5).

### Immunofluorescence

Cells were fixed for immunofluorescent probing on coverslips coated with poly-L-lysine. Cells were washed twice in PBS supplemented with 0.1 mM CaCl_2_ and 1.0 mM MgCl_2_ and fixed for 20 min in 4% paraformaldehyde (PFA), with all solutions warmed to 37°C. Following permeabilization (5 min, 0.1% Triton X-100 in PBS) and blocking (60 min, 20% FBS in PBS), cells were incubated with primary antibodies diluted in blocking solution (2 hours, see Table 1). After two washes (5 min each, 0.05% Triton X-100 in PBS), cells were incubated with secondary antibodies diluted 1:1000 in blocking solution (1 hour, see Table 1) and washed two more times. Coverslips were mounted with ProLong Glass Antifade Mountant with NucBlue (Invitrogen, P36981) on glass slides (VWR, 16004-388). Images were captured with Nikon Elements BR 5.20.02 on a Nikon Eclipse E800 microscope and DS-Qi2 camera with a 100x Apo lens (NA=1.4,WD=0.13uM) objective. Golgi area was quantified with Image J/Fiji software. Signaling was thresholded to identify the areas with GM130 signaling and area was measured with the Analyze Particles function.

### Mitochondrial isolation and blue native gel electrophoresis

As described previously (Lane *et al*., 2025b), two 150 mm dishes with cells at 80-90% confluency per condition were prepared. Following trypsinization and centrifugation of the cells, the pellet was washed with PBS. Crude mitochondria were enriched according to (Wieckowski *et al*., 2009). Briefly, cells were homogenized with 20 strokes in a Potter-Elvehjem homogenizer at 6000 rpm, 4°C in isolation buffer (225 mM mannitol, 75 mM sucrose, 0.1 mM EGTA, and 30 mM Tris-HCl, pH 7.4). Mitochondria were isolated in two centrifugation steps: 600 × g for 5 min (to remove unbroken cells and nuclei) and 7000 × g for 10 min (to pellet the mitochondria). The pellet was washed once and then mitochondria were solubilized in 1.5 M aminocaproic acid, 50 mM Bis-Tris, pH 7.0, buffer with antiproteases and either 4 g/g (detergent/protein) digitonin or DDM (n-dodecyl β-D-maltoside) to preserve or dissolve supercomplexes, respectively (Wittig *et al*., 2006; Timón-Gómez *et al*., 2020). Blue native electrophoresis in 3-12% gradient gels (Novex, BN2011BX10) (Díaz *et al*., 2009; Timón-Gómez *et al*., 2020) was used to separate proteins with 10 mg/ml ferritin (404 and 880 kDa, Sigma F4503) and BSA (66 and 132 kDa) as molecular weight standards.

### Cell survival and Synergy analysis

Survival and Synergy assays and analysis were conducted as described in Lane *et al*. (2025b). Briefly, an automated Bio-Rad cell counter (Bio-Rad, TC20, 1450102) was used to count and plate cells in 96 well plates at 5,000-10,000 cells/well. Drugs were added the following day at concentrations indicated in each figure and summarized in Table 2 for 72 hours. Cells were incubated with 10% Alamar blue (Resazurin, R&D Systems #AR002) made in fresh media was added to each well for 2 hours in the incubator. Absorbance was measured using a microplate reader (BioTek, Synergy HT; excitation at 530–570 nm and emission maximum at 580–590 nm) with Gen5 software 3.11. For each experiment, background absorbance (value of an empty well with Alamar blue and no cells) was subtracted and data were normalized to the untreated condition for each genotype to calculate percent survival. For single drug experiments, individual data points represent the average survival of duplicate or triplicate treatments for each concentration.

For Synergy survival assays with two drugs, cells treated and data was analyzed as described above, with conditions depicted in the corresponding figures and summarized in Table 2. For experiments with fetal bovine serum, values were normalized to the 10% serum condition, equivalent to the normal growth media. Individual data points represent the survival of single replicates for each concentration. Synergy calculations were performed using the ZIP score with the SynergyFinder engine https://synergyfinder.org/ (Yadav *et al*., 2015; Ianevski *et al*., 2022).

### Seahorse metabolic oximetry

Extracellular flux analysis of the Mito Stress and Glycolysis Stress Tests were performed on the Seahorse XFe96 Analyzer (Seahorse Bioscience) following manufacturer recommendations and as described previously (Lane *et al*., 2025b). The day prior to the experiment, 30,000 cells/well were seeded on Seahorse XF96 V3-PS Microplates (Agilent Technologies, 101085-004). XFe96 extracellular flux assay kit probes were incubated with the manufacturer calibration solution overnight (Agilent Technologies, 102416-100) at 37°C without CO_2_. The next day, cells were washed three times in either the Mito Stress Test Media or Glycolysis Stress Test Media. The Mito Stress Test Media consisted of Seahorse XF base media (Agilent Technologies, 102353-100) with the addition of 2 mM L-glutamine (HyClone, SH30034.01), 1 mM sodium pyruvate (Sigma, S8636), and 10 mM D-glucose (Sigma, G8769). The Glycolysis Stress Test Media contained only 2 mM L-glutamine. Cells were then incubated for 1 hour at 37°C without CO_2_ injection. During this time, flux plate probes were loaded with the appropriate 10-fold concentrated drugs for the corresponding assay and calibrated. After calibration, the Seahorse cell culture plate was equilibrated and the assay initiated. Oxygen consumption rate (OCR) and extracellular acidification rate (ECAR) were measured throughout the experiment in four read cycles (basal and one for each drug addition) consisting of a 3-minute mix cycle followed by a 3-minute read cycle. After the completion of the experiment, protein was measured and used to normalize OCR and ECAR readings for each well. Following two washes in PBS supplemented with 1 mM MgCl_2_ and 100 μM CaCl_2_, cells were lysed in Buffer A and protein concentration measured by BCA (Thermo Fisher Scientific, 23227) according to manufacturer protocol. The BCA assay absorbance was read by a BioTek Synergy HT microplate reader using Gen5 software. For data analysis of OCR and ECAR, the Seahorse Wave Software version 2.2.0.276 was used. Individual data points represent the average values of a minimum of three replicates.

For the Mito Stress Test, Seahorse injection ports were filled with oligomycin A, FCCP, and rotenone mixed with antimycin A diluted in Mito Stress Test Media (see Table 2). Non-mitochondrial respiration was determined as the lowest OCR following injection of rotenone plus antimycin A. Basal respiration was calculated from the OCR just before oligomycin injection minus the non-mitochondrial respiration. Non-mitochondrial respiration was determined as the lowest OCR following injection of rotenone plus antimycin A. ATP-dependent respiration was calculated as the difference in OCR just before oligomycin injection to the minimum OCR following oligomycin injection but before FCCP injection. Maximal respiration was calculated as the maximum OCR of the three readings following FCCP injection minus non-mitochondrial respiration. For the Glycolysis Stress Test, Seahorse injection ports were filled with 10-fold concentrated solutions of D-glucose, oligomycin, and 2-Deoxy-D-glucose (2-DG) (see Table 2 for final testing conditions and catalog numbers) in Glycolysis Stress Test. Glycolysis was calculated as the difference in ECAR between the maximum rate measurements before oligomycin injection and the last rate measurement before glucose injection. Glycolytic capacity was calculated as the difference in ECAR between the maximum rate measurement after oligomycin injection and the last rate measurement before glucose injection. Glycolytic reserve was calculated as the glycolytic capacity minus glycolysis. Non-glycolytic acidification was defined as the last rate measurement prior to glucose injection.

### Resipher

As reported in Lane *et al*. (2025b), 40,000 cells per well were seeded in 200 μL culture media in poly-L-lysine coated Nunc 96 well (Thermo, 269787) or Falcon 96 well (Falcon, 353072) plates. The next day, 150 μL of culture media was replaced by fresh, prewarmed media. The plates were covered with the Resipher sensing probe lid (Nunc Plates, NS32-N; Falcon Plates, NS32-101A) and Resipher system (Lucid Scientific, Atlanta, GA) and incubated in a humidified, 37C, 5% CO2 incubator while data was collected. After 48 hours, 10x dilutions of rotenone and antimycin A (see Table 2) in culture media were prepared and added to each well for a final concentration of 1 mM, and Resipher measurements continued for about 2 hours. Protein concentrations were determined and used to normalize OCR as described above for Seahorse plates.

### Total RNA extraction and NanoString mRNA Quantification

As described in Lane *et al*. (2025b), cells were grown on 10 cm plates to desired confluency. Cells were washed twice in ice-cold PBS containing 0.1 mM CaCl2 and 1.0 mM MgCl2. Total RNA was extracted with TRIzol (Invitrogen, 15596026) and stored at −80C. The Emory Integrated Genomics Core performed RNA extraction, RNA quality control, NanoString processing, and mRNA quantification using the Metabolic Pathways Panel (XT-CSO-HMP1-12). mRNA counts were normalized to either the housekeeping genes AARS or TBP, respectively, using NanoString nSolver software. Normalized data were further processed and visualized by Qlucore.

### ICP mass spectrometry

The protocol by Lane *et al*. (2022) was used for metal determinations, as described in Lane *et al*. (2025b). Briefly, cells were plated on 10 cm dishes until they reached desired confluency. After thoroughly washing with PBS, cells were trypsinized and pellets prepared by centrifugation (800 × g for 5 min at 4C). Samples were prepared by resuspending each pellet in ice-cold PBS, aliquoting into 3-5 tubes, and centrifugation (16,100 × g for 10 min). The final pellet was stored at −20C or −80C. Mitochondria were isolated as described above. Cell or mitochondrial pellets were digested in 50 µL 70% trace metal basis grade nitric acid (Millipore Sigma, 225711) at 95C for 10 min. After cooling, 1:40 dilutions of each sample were prepared either 2% nitric acid or 2% nitric acid with 0.5% hydrochloric acid (VWR, RC3720-16) (vol/vol). Metal levels were quantified using a triple quad ICP-MS instrument (Thermo Fisher, iCAP-TQ) operating in oxygen mode under standard conditions (RF power 1550 W, sample depth 5.0 mm, nebulizer flow 1.12L/min, spray chamber 3C, extraction lens 1,2 - 195, –-15 V). Oxygen was used as a reaction gas (0.3 mL/min) to remove polyatomic interferences or mass shift target elements (analytes measured: 32S.16O, 63Cu, 23Na, 39K, 44Ca, 56Fe, 66Zn). External calibration curves were generated using a multielemental standard (ICP-MSCAL2-1, AccuStandard, USA) and ranged from 0.5 to 1000 µg/L for each element. Scandium (10 µg/L) was used as internal standards and diluted into the sample in-line. Samples were introduced into the ICP-MS using the 2DX PrepFAST M5 autosampler (Elemental Scientific) equipped with a 250 µL loop and using the 0.25 mL precision method provided by the manufacturer. Serumnorm (Sero, Norway) was used as a standard reference material, and values for elements of interest were within 20% of the accepted value. Quantitative data analysis was conducted with Qtegra software, and values were exported to Excel for further statistical analysis.

### TMT mass spectrometry for proteomics

Cells were collected, prepared, and analyzed simultaneously with our previously reported findings as described in detail in Lane *et al*. (2025b).

### Purine nucleotide quantification

Cells were cultured on 10cm plates until 90% confluent. On a 37°C hot plate, cells were washed twice with 37°C PBS. 500 μL cold 0.1 M perchloric acid (Sigma 244252-100ML) was added directly to the cell culture dish, and the cells were scraped and transferred to a 1.5 mL tube and disrupted by probe sonication (10x 1s bursts). Homogenates were centrifuged at 10,000 × g for 10 minutes at 4°C. The supernatant was transferred to a new tube, and the pellet was saved for protein measurement. The supernatant pH was adjusted to 7 with 3.5 M K_2_CO_3_ (Oakwood Chemical 102941). Samples were stored on ice for 10-15 min to precipitate potassium perchlorate and centrifuged at 10,000 × g for 10 minutes at 4°C. To filter any remaining particulate matter, samples were centrifuged in 0.45 μm PVDF microcentrifuge filter tubes (Thermo Scientific, F2517-5) at 5000 × g for 5 minutes at 4°C. Samples were then frozen at - 80°C. For protein measurements, pellets were dissolved in 2% SDS, and 10 μL sample was diluted 1:5 in 2% SDS (total volume 50 μL). Protein was measured in duplicate by BCA as described above for Seahorse with a standard curve of 0-10 mg/mL BSA.

The day of purine nucleotide quantification, samples were thawed on ice and filtered by 0.2 μm PVDF microcentrifuge filter tubes (Thermo Scientific, F2517-6) at 5000 × g for 5 minutes at 4°C. 200 μL sample was loaded directly into HPLC loading vials (Waters, 186002639). Purine nuclotides were measured by ultra high performance liquid chromatography ACQUITY H-CLASS UPLC-PDA (Waters). Analytes were separated using reverse-phase chromatography on a Waters ACQUITY Premier HSS C18 VanGuard FIT Column, 1.8 µm, 2.1 mm x 100 mm. Elution was conducted at 0.5 mL/min with a stepped gradient of buffer A (10 mM ammonium acetate and 1.5 mM tetrabutylammonium phosphate, pH 5.0) and buffer B (10 mM ammonium phosphate, 1.5 mM TBAP, 25% acetonitrile, pH 7.0 before adding acetonitrile). The gradient consistent of the following sequence: 100% buffer A for 11 min; A linear gradient to 75% buffer B over 1 min, 3 min at 75% buffer B, a linear gradient to 100% buffer B over 4 min, 100% buffer B for 4 min, and a linear gradient to 0% buffer A over 30 sec. The column was then re-equilibrated with 100% buffer A for 9.5 min prior to next run. Purines were identified by comparing their retention times and spectral profiles to known standards, quantified at a detection wavelength of 254 nm. The injection volume was 5 μL. The column temperature was 40°C and autosampler was 4°C.

### Drosophila *survival*

All fly strains were isogenized and bred into the same genetic background and are listed in Table 3. Flies were reared at 25C on standard molasses media (Genessee Scientific) on a 12hr:12hr light:dark cycle. Male and female flies were collected at eclosion and housed separately in replicate vials of no more than 25 flies. The number of dead flies as assessed every 1-2 days and fresh food was provided every 3-4 days.

**Table 3.** *Drosophila* strains.

| Short name | Genotype | Source |
| --- | --- | --- |
| ATP7-OE | UAS-ATP7-wt-FLAG | Gift from RB (Norgate *et al.*, 2006) |
| COX17-IR | w1118; P{GD15290}v29838 | VDRC v29838 |
| Control | w1118 | BDSC 5905 |
| S6K-IR | y[1] v[1]; P{y[+t7.7] v[+t1.8]=TRiP.HMS02267}attP2 | BDSC 41702 |
| pnr-GAL4 | y[1] w[1118]; P{w[+mW.hs]=GawB}pnr[MD237]/TM3, P{w[+mC]=UAS-y.C}MC2, Ser[1] | BDSC 3039 |

Survival analysis was performed with ChatGTP 5.1 using a custom R script that reconstructs individual-level event data from percent survival and starting cohort size, followed by Kaplan–Meier estimation, log-rank testing (FDR-corrected by Benjamini-Hochberg), Cox proportional hazards modeling (Genotype × Sex interaction), hazard ratio computation, and network-based visualization of significant survival differences (Supplemental Methods). These results were confirmed against analysis of lifespan data performed in R (version 4.4.1) using RStudio (v2025.09.2 Build 418; Posit Software, PBC), also via the Kaplan-Meier method. Differences among genotypes were assessed with the log-rank test, pairwise log-rank comparisons were performed using the pairwise_survdiff() function, and p-values were adjusted for multiple comparisons as described above.

### Data availability

The mass spectrometry proteomics data will be deposited to the ProteomeXchange Consortium via the PRIDE (Perez-Riverol *et al*., 2022) partner repository.

### Bioinformatic and statistical analyses

Transcriptomics and proteomics data were processed with Qlucore Omics Explorer Version 3.6(33). Data were normalized to a variance of 1 and an average of 0 for statistical analysis and thresholding. Permutation statistical analyses were performed with the engine https://www.estimationstats.com/ with a two-sided permutation t-test using 5000 reshuffles (Ho *et al*., 2019). ANOVA and paired analyses were conducted with Prism Version 10.5.0 (774). Gene ontology studies were performed with Metascape (Zhou *et al*., 2019) and ENRICHR (Chen *et al*., 2013; Kuleshov *et al*., 2016; Xie *et al*., 2021). Kinase activity was inferred using the Kinase-Substrate Enrichment Analysis (KSEA) app (Casado *et al*., 2013; Wiredja *et al*., 2017).All violin plots depict the median, 25th, and 75th percentile values.

## Supporting information

Supplemental Methods - Code for survival

## Figure Legends

**Figure 1S.**
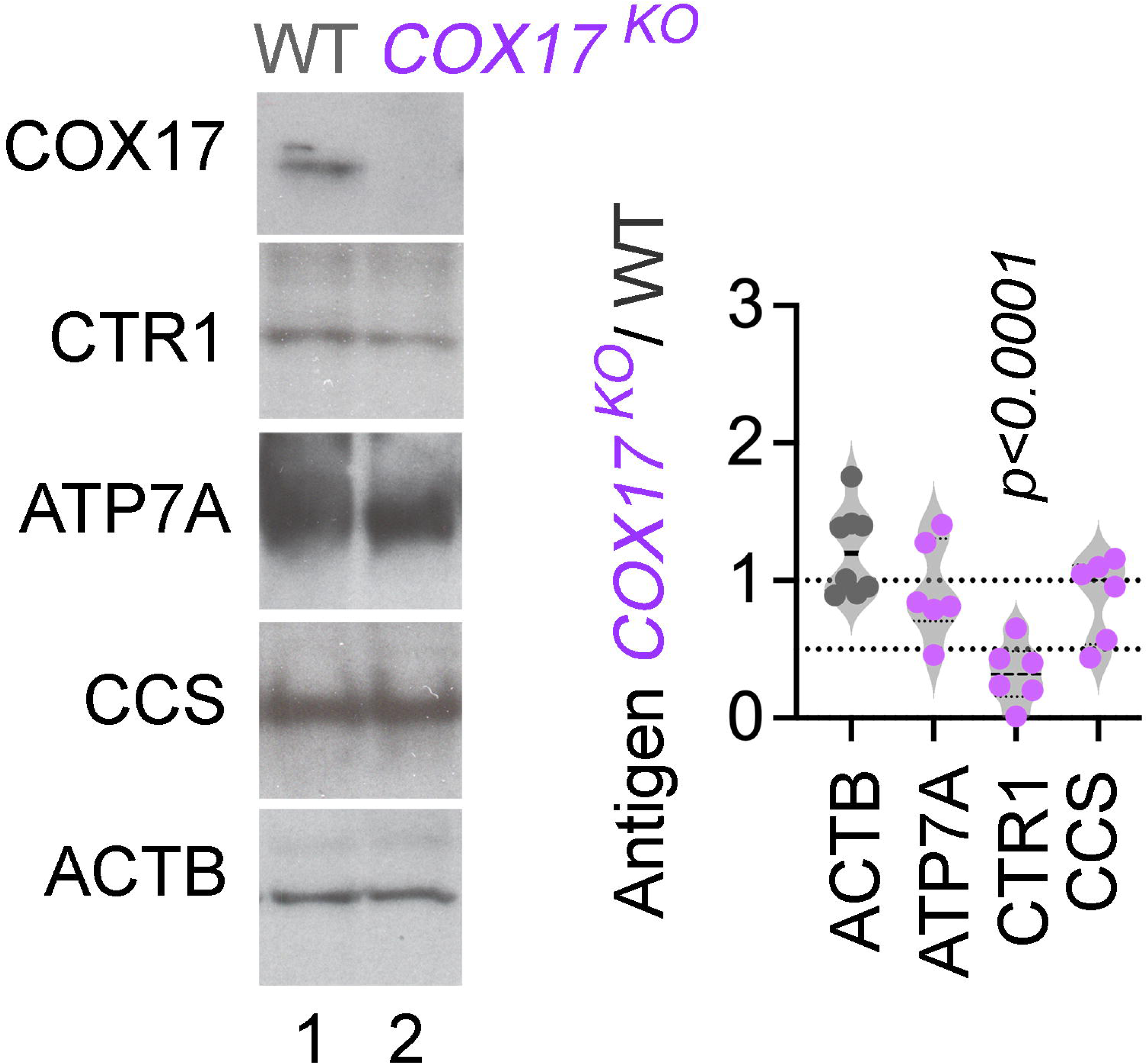
Cuproprotein levels in COX17 KO cells. Immunoblots of whole cell extracts from wild type and COX17 mutant cells probed for COX17, ATP7A, CTR1, CCS, and ACTB as a loading control. Immunoblots were quantified by normalizing protein abundance to wild-type cells. Italicized numbers represent p values analyzed by unpaired two-sided permutation t-test.

**Figure 3S1.**
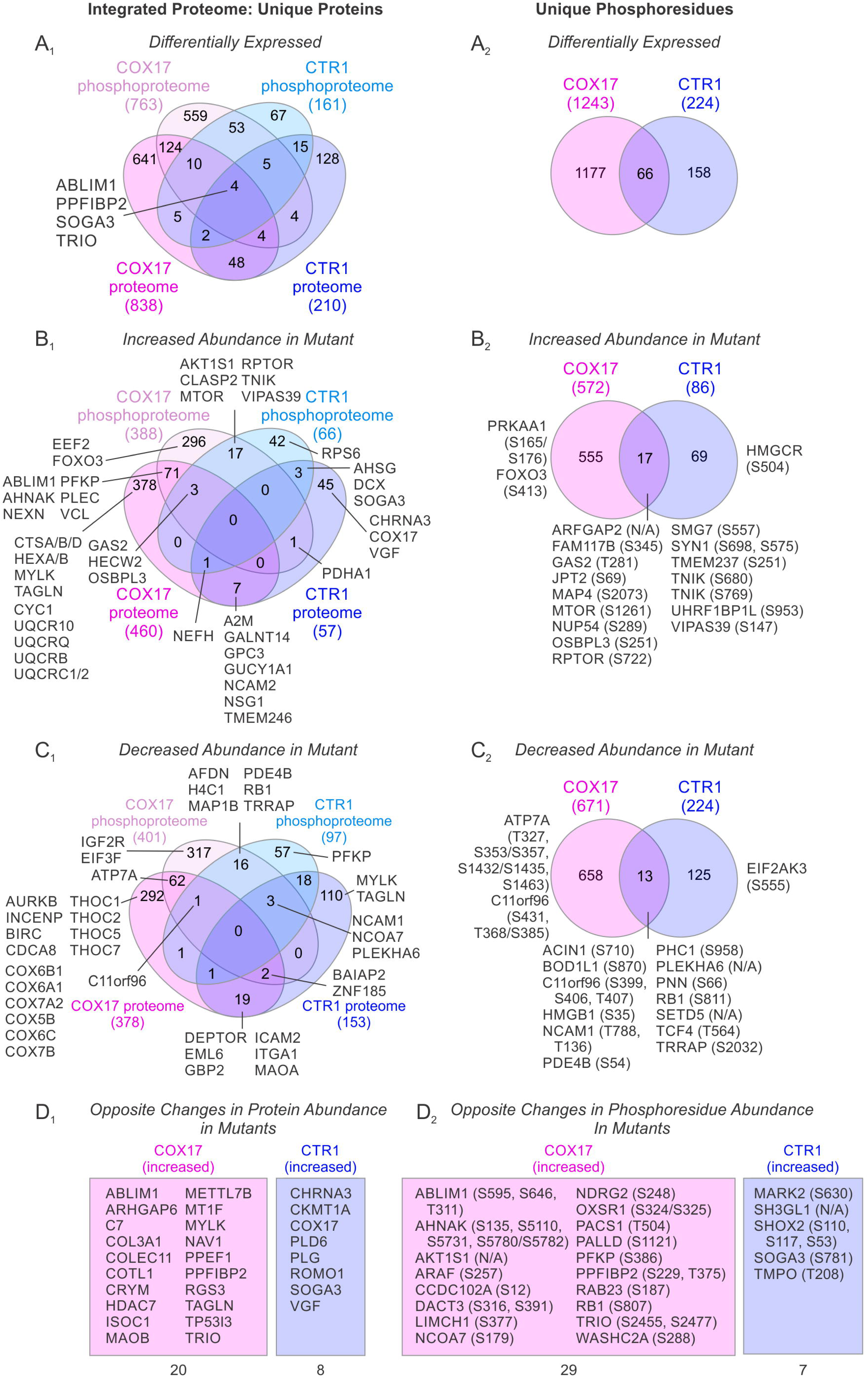
Comparative visual summary of the proteome and phosphoproteome of COX17 and CTR1 KO cells. **A-C**. Venn diagrams depicting overlap between COX17 and CTR1 KO cells of the integrated proteome (left, 1) and phosphoproteome (right, 2) that is differentially expressed (**A**), has increased abundance in mutant cells relative to wild-type cells (**B**), or has decreased abundance in mutant cells (**C**). **D**. Proteins and phosphoresidues which change in abundance in opposite directions in COX17 and CTR1 mutant cells.

**Figure 3S2.**
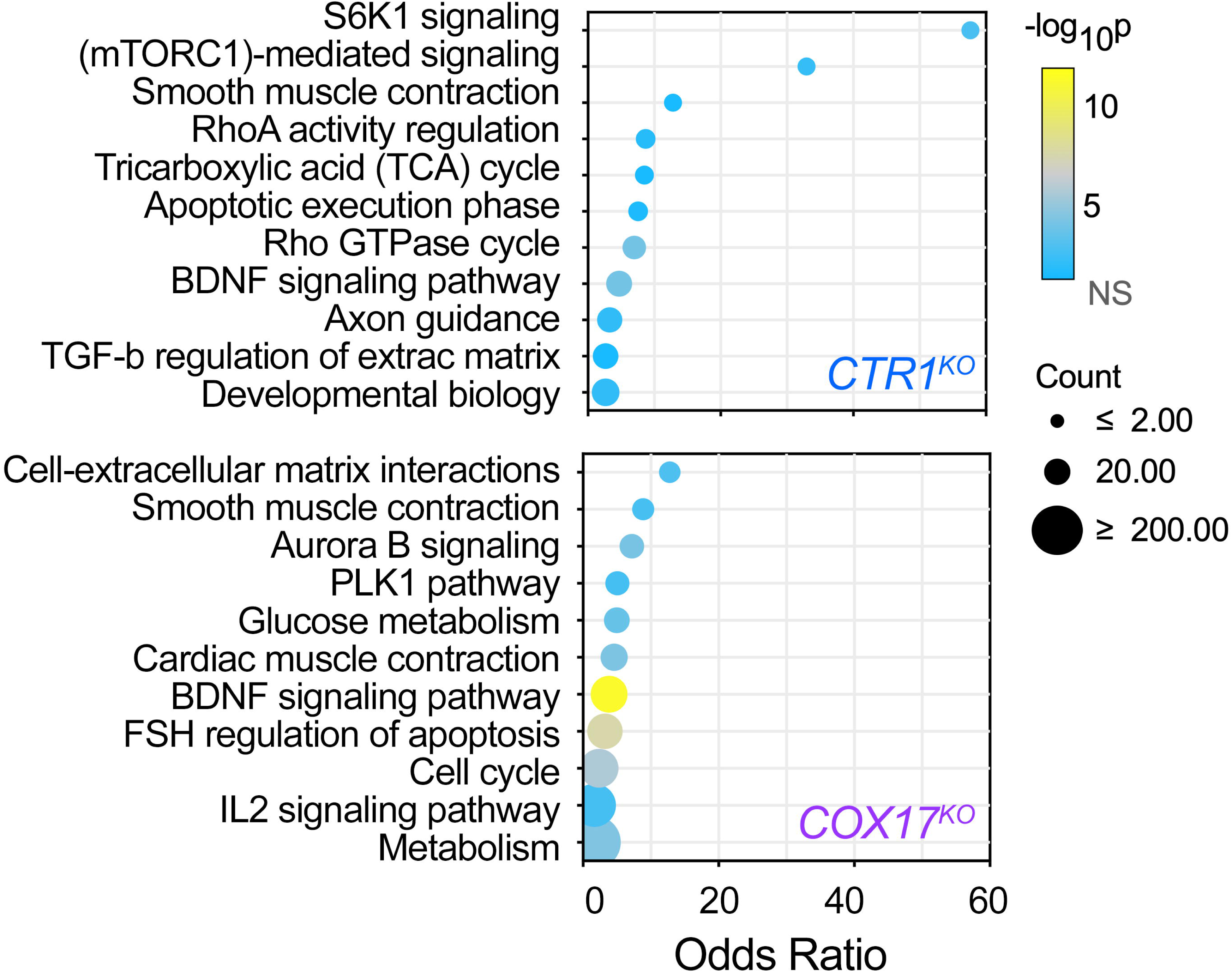
Comparative analysis of gene ontologies enriched in the COX17 and CTR1 KO proteome and phosphoproteomes. Gene ontology analysis of the merged dataset of differentially expressed proteins plus phosphopeptides in CTR1 KO and COX17 cells. Top terms based on corrected p value for each were plotted independently from each other. The BioPlanet database was queried with the ENRICHR engine (Chen *et al*., 2013; Kuleshov *et al*., 2016; Huang *et al*., 2019; Xie *et al*., 2021). Fisher exact test followed by Benjamini-Hochberg correction. CTR1 KO cell data was obtained concurrently and was reported in Lane *et al*. (2025b). See Supplemental Data File 1 for source data.

**Figure 4S.**
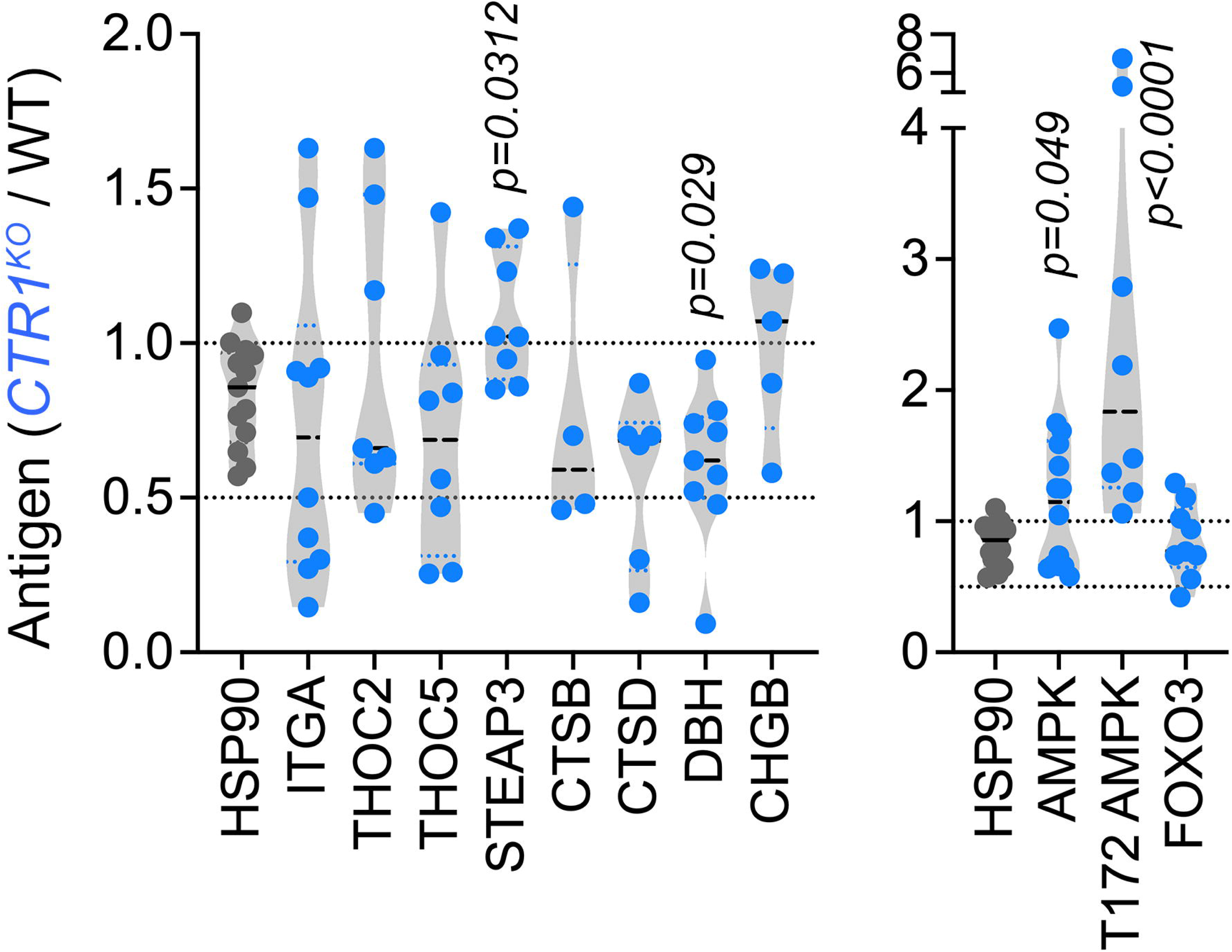
Immunoblot quantification of GO Term-selected proteins in CTR1-null cells. Quantification of immunoblots of whole cell extracts from Figure 4 (probed with antibodies for ITGA1, THOC2, THOC5, STEAP3, CTSB, CTSD, CHGB, and DBH) and Figure 6 (probed with antibodies for AMPK, AMPK pT172, and FOXO3), using HSP90 as a loading control. Immunoblots were quantified by normalizing CTR1 KO protein abundance to wild-type cells. Italicized numbers represent p values analyzed by unpaired mean difference two-sided permutation t-test.

**Figure 5S.**
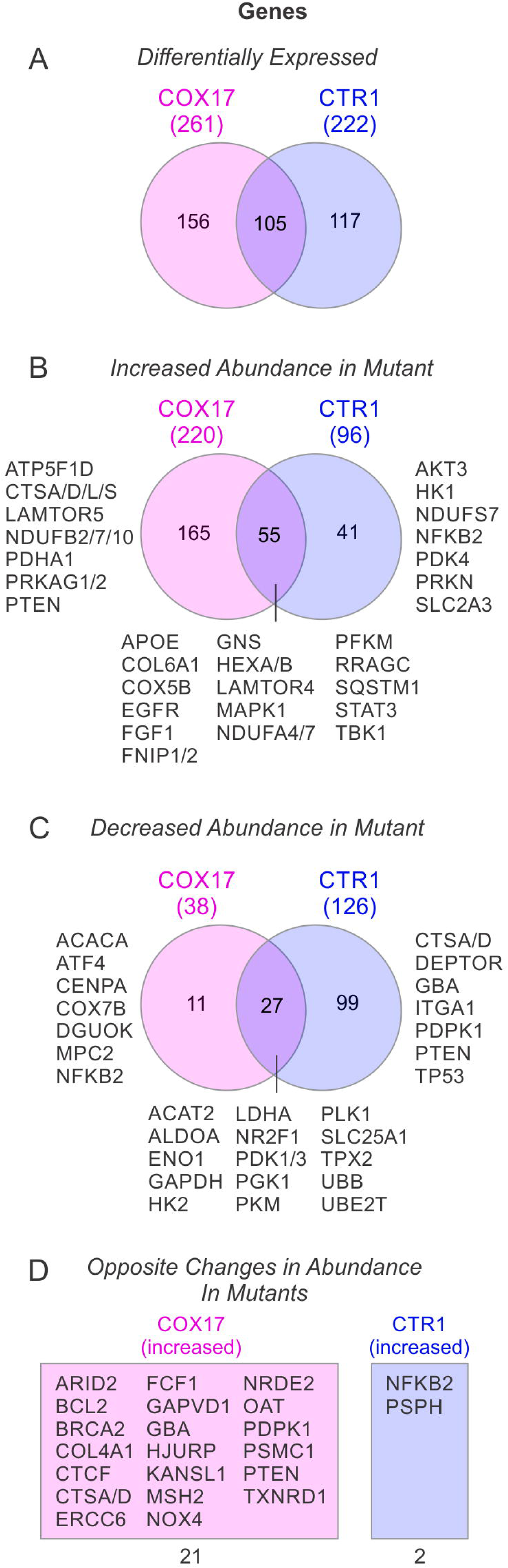
Comparative visual summary of NanoString curated metabolic transcripts in COX17 and CTR1 KO cells. **A-C**. Overlap in expression between COX17 and CTR1 KO cells of NanoString curated metabolic transcripts. Venn diagrams compare genes that are differentially expressed (**A**), have increased abundance in mutant cells relative to wild-type cells (**B**), or have decreased abundance in mutant cells (**C**). **D**. Genes that change in abundance in opposite directions in COX17 and CTR1 mutant cells.

**Figure 6S.**
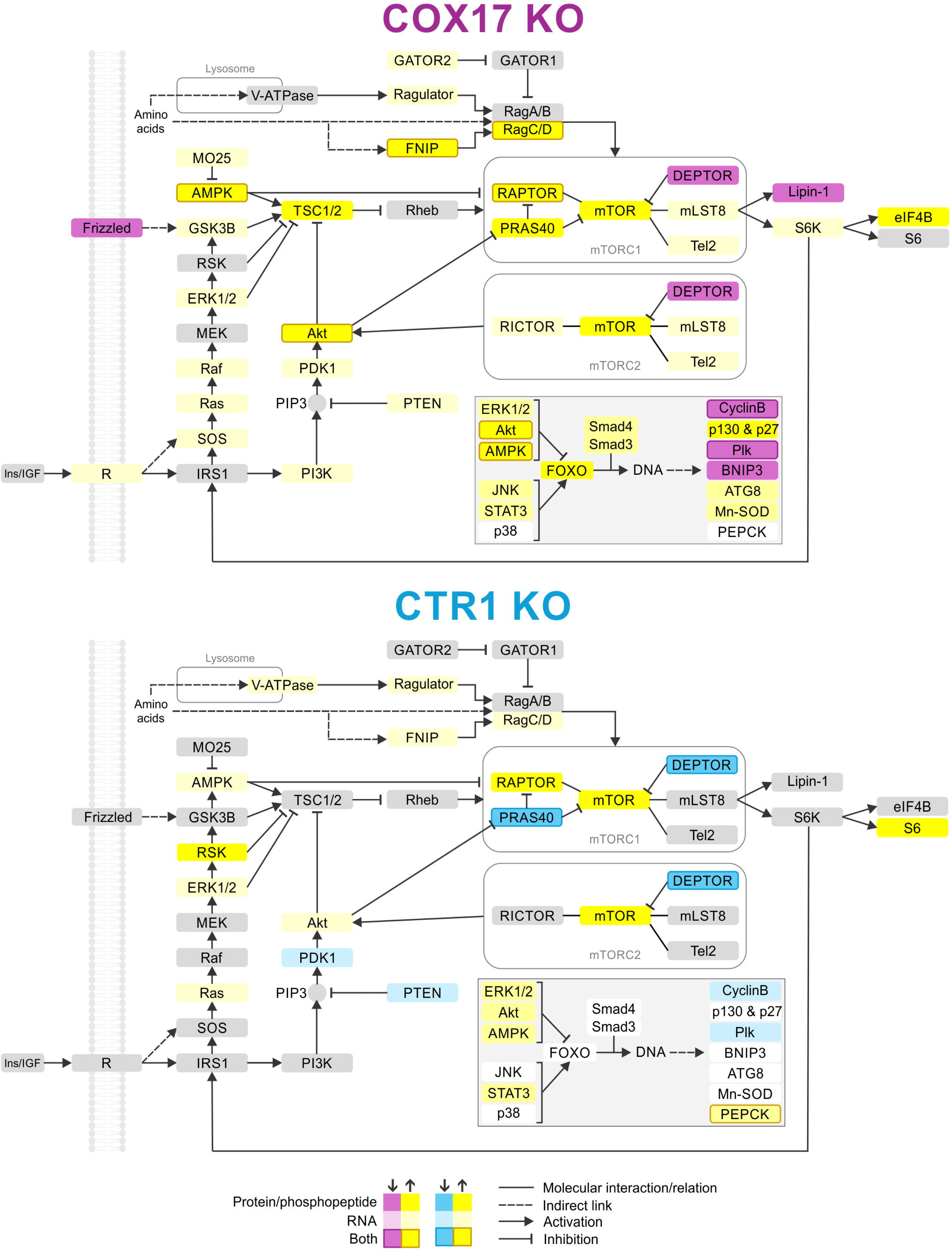
Bioinformatics summary of mTOR-AMPK pathway activity in COX17-and CTR1-null cells. Pathway diagrams summarizing abundance of RNA and proteins or phosphopeptides in the mTOR-AMPK-FOXO pathways in COX17-null (top) and CTR1-null (bottom) cells. KEGG pathway diagrams for the mTOR signaling pathway (hsa04150), AMPK signaling pathway (hsa04152), and FOXO pathway (hsa04068) were generated from the integrated proteome and NanoString RNA expression data (thresholded by q < 0.05 and fold of change ≥ 1.5) using the Color tool and were modified to illustrate all three pathways. Darker colors indicate changes at the protein or phosphopeptide level, lighter colors indicate changes at the RNA level, and borders indicate changes in both RNA and either protein or phosphopeptide levels of the target gene (see legend). Yellow indicates increased abundance and purple (COX17) or blue (CTR1) indicates decreased abundance.

**Figure S9.**
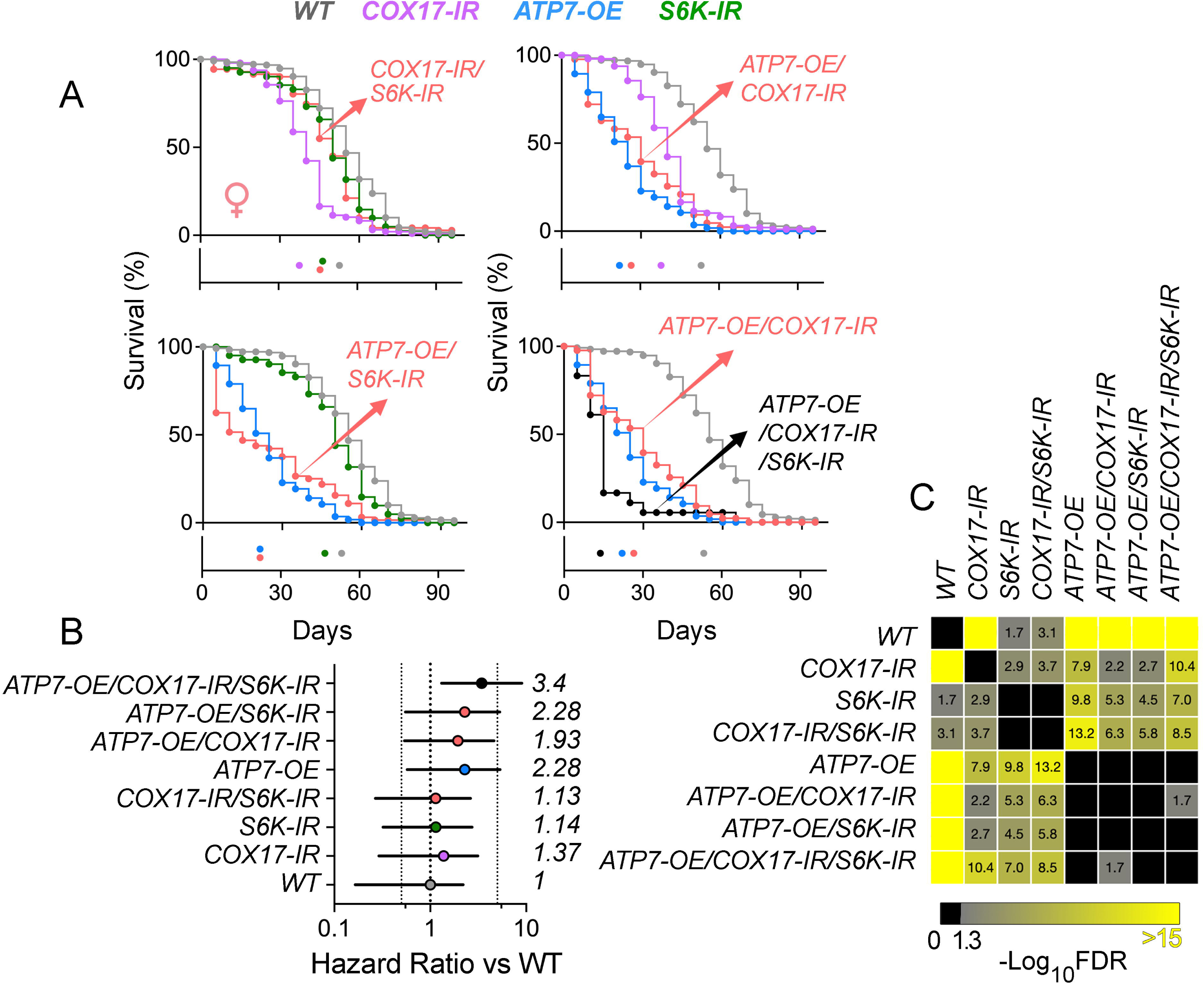
Female survival and hazard structure in copper dyshomeostasis-S6K interaction genotypes. **A.** Kaplan–Meier survival curves for females. Eight genotypes were tested: *pnr_W1118* (WT reference, n=240), *pnr_COX17_RNAi_111465* (COX17-IR, n=97), *pnr_S6K_RNAi* (S6K-IR, n=41), the combined *pnr_COX17_111465_S6K_RNAi* double knockdown (COX17-IR/SK6-IR, n=71), ATP7 overexpression (*ATP7wt_pnr_W1118, ATP7-OE, n=57*), and ATP7 overexpression combined with COX17 (n=43) and/or S6K (n=64 and 18 for triple transgenic) knockdown. Survival was reconstructed from percent-alive data and starting cohort size per genotype. Male Kaplan–Meier curves show strong survival loss following COX17 and S6K reduction, with additive or greater-than-additive decline in double and ATP7-sensitized backgrounds. **B.** Hazard ratio (HR) forest plots relative to WT. Per-genotype hazards were estimated using death rate per animal-day, normalized to WT hazard. Points represent HR, error bars indicate 95% CIs. **C.** Pairwise log-rank comparison heatmaps (Benjamini–Hochberg FDR). Matrix shows – log_10_(FDR) for all pairwise genotype survival.

## Supplementary Materials

Supplemental Data File 1 - Source Data for COX17 KO Cell Proteomics, Transcriptomics, and Gene Ontologies.xlsx

Supplemental Methods - Code for survival analysis.

## Acknowledgments

This work was supported by NIH grants 1RF1AG060285 to V.F., 1F31NS127419 and T32AG087922-02 to A.R.L., R01ES034796 to E.W, and 9R15AG074113-02 to A.V-M. A.R.L. is supported by an ARCS Foundation Award, Robert W. Woodruff Fellowship, and John B. Lyon Memorial Scholarship Award. This study was supported in part by the Emory Integrated Genomics and Proteomics Cores, which are subsidized by the Emory University School of Medicine, and the Emory HPLC Bioanalytical Core (RRID:SCR_023531). V.F. is grateful for mitochondria provided by Maria Olga Gonzalez.

