## Supplemental Methods - Code for survival for "Distinct signaling mechanisms and proteome phenotypes are elicited by compartment-specific genetic defects of copper homeostasis": README_survival_pipeline.md

# Survival Analysis Pipeline (Genotype × Sex)

This repository contains a ready-to-use R script for Kaplan–Meier survival analysis, sex-specific comparisons, and genotype interaction modeling.

The pipeline is designed for experiments with:
- Multiple **genotypes**
- **Male and female** cohorts
- Survival recorded over time as **percent survival**

---

## Files

- `survival_analysis_pipeline.R`  
  Main analysis script. Reconstructs individual events from % data + N₀ and performs:
  - Kaplan–Meier curve estimation per group
  - Mean survival time calculation
  - Log-rank tests (Male vs Female within each genotype)
  - Pairwise log-rank tests between all genotypes per sex
  - Cox proportional hazards model with Sex × Genotype interaction
  - Hazard ratios vs WT per sex (with 95% CI)
  - Heatmaps of BH-adjusted p-values
  - Network of significant survival differences (p < 0.05)

- `survival_percent_example.csv`  
  Minimal example dataset of percent survival over time for **three genotypes** and both sexes.

- `N0_by_sex_example.csv`  
  Minimal example of starting animal numbers for each genotype and sex.

---

## Input File Formats

### 1. `survival_percent.csv`

Your real data file **must** follow this structure:

- First column: `days` (time points, in days or other consistent units)
- Remaining columns: percent survival values (0–100) for each genotype × sex.

Example (wide) format:

```csv
days,pnr_W1118 m,pnr_COX17_RNAi_111465 m,pnr_S6K_RNAi m,pnr_W1118 f,pnr_COX17_RNAi_111465 f,pnr_S6K_RNAi f
0,100,100,100,100,100,100
10,95,90,92,98,96,94
20,80,60,70,90,85,82
30,50,30,40,70,60,55
40,10,5,8,30,20,15
```

Column naming convention:
- A **space** followed by `m` or `f` indicates male or female,
  e.g., `pnr_W1118 m`, `pnr_W1118 f`.
- The text **before** the space is the **base genotype name**, which
  must match the `Group` column in `N0_by_sex.csv`.

### 2. `N0_by_sex.csv`

This file specifies starting animal counts for each genotype and sex.

Required columns:
- `Group` — base genotype name (must match the portion before " m" / " f" in `survival_percent.csv`).
- `m` — initial number of males.
- `f` — initial number of females.

Example:

```csv
Group,m,f
pnr_W1118,50,60
pnr_COX17_RNAi_111465,40,45
pnr_S6K_RNAi,35,38
```

---

## Running the Pipeline

1. Place the following files in the same folder:
   - `survival_analysis_pipeline.R`
   - Your experimental `survival_percent.csv`
   - Your experimental `N0_by_sex.csv`

   For testing, you can use the provided example files:
   - `survival_percent_example.csv` → copy/rename to `survival_percent.csv`
   - `N0_by_sex_example.csv` → copy/rename to `N0_by_sex.csv`

2. Open R and set the working directory:

```r
setwd("path/to/your/folder")
source("survival_analysis_pipeline.R")
```

3. The script will create an `output/` directory containing:
   - `KM_event_data.csv` — reconstructed subject-level survival data
   - `KM_curves_Male.pdf`, `KM_curves_Female.pdf`
   - `Mean_survival_time_by_group.csv`
   - `Sex_logrank_by_genotype.csv`
   - `Pairwise_logrank_Males_BH.csv`, `Pairwise_logrank_Females_BH.csv`
   - `Cox_Sex_by_Genotype_interaction.csv`
   - `HazardRatios_Males.csv`, `HazardRatios_Females.csv`
   - `Heatmap_logrank_Males.png`, `Heatmap_logrank_Females.png`
   - `Significant_Survival_Network_Edges.csv`

---

## Adapting to Your Experiment

- Add or remove genotypes by adding/removing columns in `survival_percent.csv` and rows in `N0_by_sex.csv`.
- Timepoints can be irregular (e.g., 0, 5, 7, 10 days); the script uses them as provided.
- You may change the "wild-type" reference genotype for hazard ratios by editing the `wt_m` and `wt_f` definitions near the bottom of `survival_analysis_pipeline.R`.

---

## Citation / Acknowledgment

If you share this pipeline or use it in a publication, you may describe it as:

> “Survival analysis was performed using a custom R script that reconstructs individual-level event data from percent survival and starting cohort size, followed by Kaplan–Meier estimation, log-rank testing, Cox proportional hazards modeling (Genotype × Sex interaction), hazard ratio computation, and network-based visualization of significant survival differences.”
