## Supplemental Methods - Code for survival for "Distinct signaling mechanisms and proteome phenotypes are elicited by compartment-specific genetic defects of copper homeostasis": survival_analysis_pipeline.R

```
############################################################
# Survival Analysis Pipeline for Genotype × Sex Experiments
# Author: YOUR NAME
# Description:
#   - Input:  (1) survival_percent.csv  (2) N0_by_sex.csv
#   - Output: KM curves, hazard ratios, log-rank statistics,
#             sex × genotype Cox model, heatmaps & network.
#   - Ready for reuse in manuscripts and shared analysis.
############################################################

## ======================= USER GUIDE =======================
## Required input files in the working directory:
##
## 1) survival_percent.csv
##    columns:
##      days, g1_m, g2_m, ..., gK_m, g1_f, g2_f, ..., gK_f
##    where each column (except 'days') is % survival (0–100).
##
## 2) N0_by_sex.csv
##    columns:
##      Group, m, f
##    where Group matches the base genotype name used in
##    column headers (before the space and sex suffix).
##
## Example:
##   Group,m,f
##   pnr_W1118,187,240
##   pnr_COX17_RNAi_111465,100,97
##
## To run:
##   setwd("path/to/your/data")
##   source("survival_analysis_pipeline.R")
##
## All outputs are written into ./output
## ==========================================================

suppressPackageStartupMessages({
  library(tidyverse)
  library(survival)
  library(pheatmap)
  library(broom)
})

dir_out <- "output"
if (!dir.exists(dir_out)) dir.create(dir_out)

message(">>> Loading input files")

surv_raw <- read.csv("survival_percent.csv", check.names = FALSE)
N0_tab   <- read.csv("N0_by_sex.csv", check.names = FALSE)

# helper: extract base genotype & sex from column name
extract_base <- function(colname){
  # remove trailing " m" or " f"
  gsub(" m$| f$", "", colname)
}
extract_sex <- function(colname){
  ifelse(grepl(" f$", colname), "F",
         ifelse(grepl(" m$", colname), "M", NA))
}

############################################################
# 1. Convert % → survivors → deaths → KM event table
############################################################
message(">>> Reconstructing events from % survival and N0")

# Build long table: days, group, percent, sex, base, N0
long <- surv_raw %>%
  pivot_longer(-days, names_to = "group", values_to = "percent") %>%
  mutate(
    sex  = extract_sex(group),
    base = extract_base(group)
  )

# join N0
long <- long %>%
  left_join(N0_tab, by = c("base" = "Group")) %>%
  mutate(
    N0 = ifelse(sex == "M", m, f)
  )

if (any(is.na(long$N0))) {
  stop("Some groups have missing N0. Check N0_by_sex.csv naming.")
}

# reconstruct survivors and deaths per group
long <- long %>%
  group_by(group) %>%
  arrange(days) %>%
  mutate(
    survivors = round(percent/100 * N0),
    deaths    = pmax(lag(survivors, default = N0) - survivors, 0)
  ) %>%
  ungroup()

# event records
events <- long %>%
  filter(deaths > 0) %>%
  uncount(deaths) %>%
  transmute(
    time   = days,
    status = 1,
    group,
    sex,
    base
  )

# censored at last time for remaining survivors
last_day <- max(long$days)
censored <- long %>%
  group_by(group) %>% slice_tail(n = 1) %>%
  filter(survivors > 0) %>%
  uncount(survivors) %>%
  transmute(
    time   = last_day,
    status = 0,
    group,
    sex,
    base
  )

km_data <- bind_rows(events, censored)

write.csv(km_data,
          file.path(dir_out, "KM_event_data.csv"),
          row.names = FALSE)

############################################################
# 2. Kaplan–Meier curves by sex
############################################################
message(">>> Fitting Kaplan–Meier curves")

km_data$group <- factor(km_data$group)
km_data$sex   <- factor(km_data$sex, levels = c("M","F"))

make_km_plot <- function(sex_label){
  sub <- subset(km_data, sex == sex_label)
  fit <- survfit(Surv(time, status) ~ group, data = sub)
  pdf(file.path(dir_out,
                paste0("KM_curves_", ifelse(sex_label=="M","Male","Female"), ".pdf")),
      width = 7, height = 5)
  plot(fit, col = 1:length(fit$strata), lwd = 2,
       xlab = "Time (days)", ylab = "Survival probability",
       main = paste("Kaplan–Meier:", ifelse(sex_label=="M","Males","Females")))
  legend("topright", legend = names(fit$strata),
         col = 1:length(fit$strata), lwd = 2, cex = 0.6)
  dev.off()
}

make_km_plot("M")
make_km_plot("F")

############################################################
# 3. Mean survival time per group
############################################################
message(">>> Computing mean survival time via KM")

mean_survival <- km_data %>%
  group_by(group) %>%
  do({
    fit <- survfit(Surv(time, status) ~ 1, data = .)
    data.frame(mean_time = summary(fit, times = NULL)$table["*rmean"])
  }) %>%
  ungroup()

write.csv(mean_survival,
          file.path(dir_out, "Mean_survival_time_by_group.csv"),
          row.names = FALSE)

############################################################
# 4. Log-rank tests: Male vs Female within each genotype
############################################################
message(">>> Running log-rank tests: sex differences per genotype")

bases <- unique(km_data$base)
sex_comp <- data.frame(
  Genotype = bases,
  P_value  = NA_real_
)

for(i in seq_along(bases)){
  g <- bases[i]
  sub <- subset(km_data, base == g)
  if (length(unique(sub$sex)) < 2) next
  lr   <- survdiff(Surv(time, status) ~ sex, data = sub)
  pval <- 1 - pchisq(lr$chisq, df = 1)
  sex_comp$P_value[i] <- pval
}

sex_comp$FDR_BH <- p.adjust(sex_comp$P_value, method = "BH")
sex_comp <- arrange(sex_comp, FDR_BH)

write.csv(sex_comp,
          file.path(dir_out, "Sex_logrank_by_genotype.csv"),
          row.names = FALSE)

############################################################
# 5. Pairwise log-rank tests within sex (all genotypes)
############################################################
message(">>> Running pairwise log-rank tests within each sex")

do_pairwise <- function(sex_label){
  sub <- subset(km_data, sex == sex_label)
  pw  <- pairwise_survdiff(Surv(time, status) ~ group,
                           data = sub, p.adjust.method = "BH")
  mat <- pw$p.value
  mat[is.na(mat)] <- 1
  return(mat)
}

pm <- do_pairwise("M")
pf <- do_pairwise("F")

write.csv(pm, file.path(dir_out,"Pairwise_logrank_Males_BH.csv"))
write.csv(pf, file.path(dir_out,"Pairwise_logrank_Females_BH.csv"))

############################################################
# 6. Cox model with Sex × Genotype interaction
############################################################
message(">>> Fitting Cox model with Sex × Genotype interaction")

km_data$base <- factor(km_data$base)
cox_int <- coxph(Surv(time, status) ~ base * sex, data = km_data)
summary_int <- broom::tidy(cox_int, exponentiate = TRUE, conf.int = TRUE)

write.csv(summary_int,
          file.path(dir_out, "Cox_Sex_by_Genotype_interaction.csv"),
          row.names = FALSE)

############################################################
# 7. Hazard ratios vs WT for each sex
############################################################
message(">>> Computing hazard ratios vs WT per sex")

# choose WT as first base name (can be edited by user)
wt_m <- paste0(bases[1], " m")
wt_f <- paste0(bases[1], " f")

compute_HR <- function(sex_label, wt_group){
  sub <- subset(km_data, sex == sex_label)
  groups <- levels(sub$group)
  res <- lapply(groups, function(g){
    df <- subset(sub, group == g)
    D  <- sum(df$status == 1)
    Tt <- sum(df$time)
    lam <- D / Tt
    data.frame(group = g, deaths = D, time = Tt, hazard = lam)
  })
  hr <- bind_rows(res)
  lam0 <- hr$hazard[hr$group == wt_group]
  D0   <- hr$deaths[hr$group == wt_group]
  hr <- hr %>%
    mutate(
      HR = hazard / lam0,
      var_logHR = 1/deaths + 1/D0,
      se_logHR  = sqrt(var_logHR),
      HR_low  = exp(log(HR) - 1.96*se_logHR),
      HR_high = exp(log(HR) + 1.96*se_logHR)
    )
  hr
}

hr_m <- compute_HR("M", wt_m)
hr_f <- compute_HR("F", wt_f)

write.csv(hr_m, file.path(dir_out,"HazardRatios_Males.csv"), row.names = FALSE)
write.csv(hr_f, file.path(dir_out,"HazardRatios_Females.csv"), row.names = FALSE)

############################################################
# 8. Heatmaps of BH-adjusted pairwise p-values
############################################################
message(">>> Generating log-rank heatmaps")

pheatmap(pm,
         main = "Males: BH-adjusted log-rank p-values",
         color = colorRampPalette(c("yellow","red","black"))(100),
         filename = file.path(dir_out,"Heatmap_logrank_Males.png"))

pheatmap(pf,
         main = "Females: BH-adjusted log-rank p-values",
         color = colorRampPalette(c("yellow","red","black"))(100),
         filename = file.path(dir_out,"Heatmap_logrank_Females.png"))

############################################################
# 9. Network of significant genotype differences (p < 0.05)
############################################################
message(">>> Building significant survival difference network (p < 0.05)")

build_edge_table <- function(mat, sex_label, alpha = 0.05){
  groups <- rownames(mat)
  edges <- list()
  for(i in seq_along(groups)){
    for(j in seq_along(groups)){
      if (j <= i) next
      p <- mat[i,j]
      if (!is.na(p) && p < alpha){
        edges[[length(edges)+1]] <- data.frame(
          Sex    = sex_label,
          Node1  = groups[i],
          Node2  = groups[j],
          p_value = p
        )
      }
    }
  }
  if (length(edges)==0) return(data.frame())
  bind_rows(edges)
}

edges_m <- build_edge_table(pm,"Male")
edges_f <- build_edge_table(pf,"Female")
edges_all <- bind_rows(edges_m, edges_f)

write.csv(edges_all,
          file.path(dir_out,"Significant_Survival_Network_Edges.csv"),
          row.names = FALSE)

message(">>> Analysis complete. All outputs written to: ", dir_out)
```
